# Non-REM sleep in major depressive disorder

**DOI:** 10.1101/2021.03.19.436132

**Authors:** Leonore Bovy, Frederik D. Weber, Indira Tendolkar, Guillén Fernández, Michael Czisch, Axel Steiger, Marcel Zeising, Martin Dresler

## Abstract

Disturbed sleep is a key symptom in major depressive disorder (MDD). REM sleep alterations are well described in the current fliterature, but little is known about non-REM sleep alterations. Additionally, sleep disturbances relate to a variety of cognitive symptoms in MDD, but which features of non-REM sleep EEG contribute to this, remains unknown. We comprehensively analyzed non-REM sleep EEG features in three independently collected datasets (*N*=284). These included MDD patients with a broad age range, varying duration and severity of depression, unmedicated or medicated, age- and gender-matched to healthy controls. We explored changes in sleep architecture including sleep stages and cycles, spectral power, sleep spindles, slow waves (SW), and SW-spindle coupling. Next, we analyzed the association of these sleep features with acute measures of depression severity and overnight consolidation of procedural memory. Overall, no major systematic alterations in non-REM sleep architecture were found in patients compared to controls. For the microstructure of non-REM sleep, we observed a higher spindle amplitude in unmedicated patients compared to controls, and after the start of antidepressant medication longer SWs with lower amplitude and a more dispersed SW-spindle coupling. In addition, long-term, but not short-term medication seemed to lower spindle density. Overnight procedural memory consolidation was impaired in medicated patients and associated with lower sleep spindle density. Our results suggest that alterations in non-REM sleep EEG might be more subtle than previously reported. We discuss these findings in the context of antidepressant medication intake and age.

**Statement of Significance:** Depression affects large and diverse populations worldwide, including their sleep. Most sleep is non-REM sleep, which is vital to cognitive function, including memory. How non-REM is affected during a depression or medical treatment remains poorly investigated. We classified non-REM sleep of depressive patients against healthy controls in unprecedented analysis detail and confidence using the largest dataset published so far while also test sleep alterations associations with impaired memory. Surprisingly, severe depression alone did not alter sleep. We observed severe non-REM sleep alterations only worsening under patient medication, which ultimately coincided with 24-hour memory impairments. Though causal influences of medication on sleep in depressive patients remains to be investigated, this cautions common clinical practice in long-term treatment with antidepressants.

## Introduction

Major depressive disorder (MDD) is a common psychiatric disorder and a serious public health problem (World Health Organization, 2017). MDD patients suffer from several physical symptoms, including subjective sleep complaints such as sleeplessness (i.e. insomnia; [1]. Objective changes in sleep quality, such as abnormalities in the efficiency and duration of sleep can also be observed [2, 3]. In addition, changes in sleep architecture and in particular rapid eye-movements (REM) sleep have been reported: increased REM density and duration, as well as decreased latency to REM sleep onset (e.g. [4]). In non-REM sleep, several studies report a decrease in the amount of slow wave sleep (SWS) in MDD patients compared to controls (as reviewed by [5]), as well as reduced slow wave activity (SWA; [6]), however see [7, 8] for non-significant findings or even increases in SWA in females only [9].

Non-REM investigations of more detailed sleep microstructures in MDD are scarce and mainly investigated sleep spindles and slow waves (SWs) with conflicting outcomes. Sleep spindles are a hallmark in electrophysiological activity defining non-REM sleep. They consist of short (0.5–2 s) waxing and waning bursts of oscillatory activity (12–15 Hz) that consistently appear throughout non-REM sleep. SWs are large (> 75 μV) waves occurring isolated but largely in the deeper stages of non-REM sleep and describe the events in the slow-wave (0.5–4 Hz), slow oscillation/lower delta (0.5–1 Hz), or upper delta (1–4 Hz) band with definitions differing across studies. Early sleep studies indicated decreases in sleep spindle activity in MDD patients [10, 11], confirmed also in high-risk groups [12] and those with manic-depression [13], but see also [8, 14] for reports on spindle features remaining either unaltered or differing dependent on sex. Notably, several limitations are present in these few studies, including limited sample sizes and mild MDD severity. Another aspect of concern is that the patients are often medicated with drugs that likely influence sleep architecture and non-REM features such as sleep spindles [15], and a detailed description of medication types is often lacking. Besides, these studies also neglect that sleep spindles and SWs do not always appear in isolation: sleep spindles couple to SWs and hippocampal sharp wave-ripples and this fine-tuned interaction is indicative of processes critical to sleep-associated memory consolidation (see [16] for a review). Despite this interplay of spindles and SWs being critical to cognitive function, to the best of our knowledge, such SW-spindle coupling has not been investigated in MDD before, and a systematic overview of all common non-REM sleep microstructure features in larger MDD samples is missing.

Next to sleep disturbances, MDD is also characterized by several cognitive deficits, including memory impairments [17]. It remains unclear how the disrupted sleep relates to the development and/or maintenance of some of these cognitive deficits. Indeed, deficits in overnight procedural memory consolidation were observed in medicated MDD patients, compared to healthy controls, with initial learning performance itself unimpaired [18, 19]. Furthermore, hippocampal activity has been linked to sleep-related procedural memory consolidation [20] and depressed patients, especially those with recurring episodes and early-onset symptomology, have been shown to have smaller hippocampal volumes compared to healthy controls [21]. In addition, previous studies have also consistently shown changes in thalamic function and structure in MDD patients, such as a reduction in grey matter [22, 23], deficits in thalamo-cortical connectivity [24], and associations between reduction in size and increased symptom severity [25]. Interestingly, a recent study reported that patients with hippocampal damage show a reduced amount of SWS as well as SWA and within the SW cycle, a delay of co-occurring sleep spindles [26].

Overall, given these memory deficits, subcortical anatomical alterations, and sleep disturbances in MDD, it seems plausible that the procedural memory deficits seen in MDD may be directly linked to particular changes in specific non-REM sleep properties, next to already reported general changes in sleep architecture, such as sleep spindle alterations or their interplay with other brain rhythms like SWs and hippocampal ripples.

In the current study, we aimed to provide a characterization of non-REM sleep in MDD by exploring systematic changes on both a macro level, including sleep architecture and power spectra but also, on a micro-level, including sleep spindles, SW and SW-spindle coupling, in the largest sample to date. Next, we investigated the influence of these sleep alterations on overnight procedural memory consolidation. Lastly, we explored how far these non-REM features were related to depression severity and outcome. We performed an identical analysis on all independently collected datasets to explore the robustness, replicability, and generalizability of our findings.

## Methods and Materials

### Participants

Three datasets were independently collected and denoted Dataset A (fully reanalyzed from partly published data in [27], here including partially overlapping and additional participants), Dataset B (unpublished), and Dataset C (partly published in [28]), of which previous sleep-targeted analyses were limited to the description of changes in the composition of scored sleep stages. Dataset A and B included each 40 MDD patients and 40 healthy controls, whereas Dataset C included 30 MDD patients and 28 healthy controls. Data were collected at the Max Planck Institute of Psychiatry in Munich, Germany.

MDD patients were matched by age and gender to controls: Dataset A was balanced on a group level whereas in Dataset B and C they were matched by age (±2 year of tolerance) and gender individually. Patient profiles between the datasets differed in age, the severity of depression at baseline, and depressive episodes (excluding the current episode, see Table 1).

**Table 1.** Demographics table of the final datasets used for polysomnography analyses (mean ± SE)

|  | Dataset A |  | Dataset B |  | Dataset C |  |
| --- | --- | --- | --- | --- | --- | --- |
|  | Controls | Patients | Controls | Patients | Controls | Patients |
| n | 40 | 40 | 40 | 38 | 28 | 30 |
| Age | 46.8 ± 1.69 | 50.1 ± 1.37 | 31.5 ± 1.62 | 31.3 ± 1.65 | 45.5 ± 3.11 | 45.6 ± 3.07 |
| Gender (male/female) | 20/20 | 19/21 | 19/21 | 18/20 | 10/18 | 11/19 |
| HAMD at baseline | NA | 24.5 ± 0.96 | NA | 19.9 ± 0.62 | NA | 25.2 ± 1.11 |
| Number prev. episodes | NA | 2.1 ± 0.34 | NA | 1.37 ± 0.25 | NA | 2.7 ± 0.56 |

Datasets A and C show a similar distribution of age, depression severity (moderate to strong), and the number of previous depressive episodes (several, typically not the first). Dataset B had a younger age distribution with less severe depression severity (moderate) and a lower number of previous depressive episodes (many patients with their first).

Polysomnography of MDD patients was recorded at two timepoints in Dataset B and C and all MDD patients received antidepressant medication eventually (see Table S7 medication types per patient). Ambidextrous people were excluded from the samples that included procedural memory tasks. Due to technical failure in the EEG data of two medicated MDD patients in Dataset B, all paired analyses were matched on the remaining full datasets (*n* = 38 per group).

### Experimental procedures and memory task

Experimental procedures were as described in [27] for Dataset A and similarly applied in Dataset B and Dataset C. Both Dataset A and B employed a common procedural memory paradigm (finger tapping task). Procedural memory data was available for all Datasets A [18, 27] and most of Dataset B participants (see Figure 1 and Table 3 for details). Dataset B in addition included a declarative memory paradigm (word-pair learning task; which was not included in the current analysis), as well as high-density EEG recordings and anatomical MRIs. Dataset B patients were unmedicated during their initial session but thereafter received medication that continued into the 1-week follow-up session. This follow-up session included the same task with a new tapping sequence and another high-density EEG recording. Dataset C patients were medicated for 7 days before the first recordings and continued for a follow-up at 28 days of medication. Thus Dataset A was mainly medicated long-term (although medication history was not consistently assessed), Dataset B short-term (i.e. 7 days), and Dataset C short- (7 days) and long-term (28 days).

**Figure 1:**
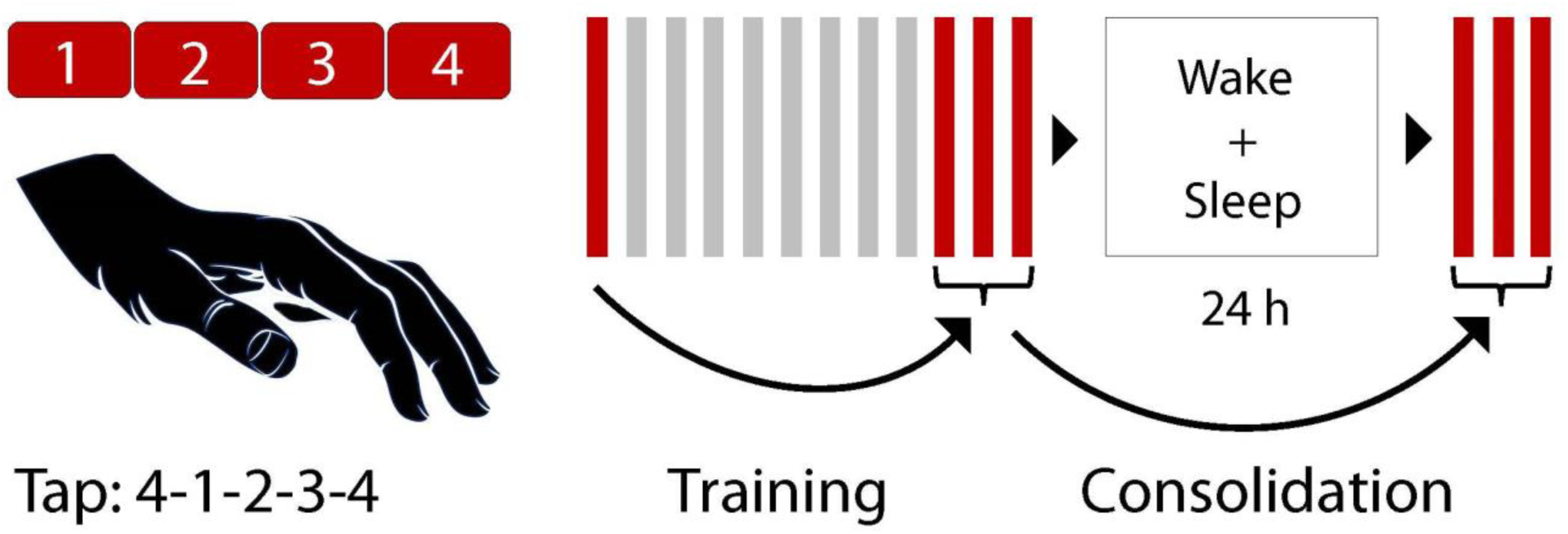
Finger tapping task design. Participants were instructed to tap a sequence (e.g. 4-1-3-2-4) as accurate and fast as possible on a computer keyboard with their non-dominant hand. The number of correct sequences was the main behavioral outcome measure. During a training session, participants had to tap the sequence for 12 runs of 30 seconds each, with a 20-second break between runs. The difference between the first run and the mean of the last three runs was considered the “training effect”. After 24 hours, which included a full night of EEG recorded sleep, three additional runs were tested. The difference between the mean of these three test runs and the mean of the last three training runs was considered the “consolidation effect”.

**Table 3.** Procedural memory results from finger tapping task (mean ± SE)

|  | Dataset A |  | Dataset B |  |  |
| --- | --- | --- | --- | --- | --- |
|  | Controls | Medicated | Controls | Unmedicated | Medicated |
| Baseline | **6.65 ± 0.576/**<br>**6.33 ± 0.493 (1rm)** | **4.15 ± 0.463\*\*** | 7.8 ± 0.76/<br>7.26 ± 0.545 (1rm) | 6.08 ± 0.606 | 7.33 ± 0.813 |
| Training | 1.75 ± 0.237/<br>1.62 ± 0.208 (1rm) | 1.76 ± 0.175 | 1.72 ± 0.381/<br>1.26 ± 0.201 (2rm) | 1.24 ± 0.353/<br>0.913 ± 0.137 (1rm) | 0.877 ± 0.247 /<br>0.654 ± 0.11 (1rm) |
| Consolidation | **0.088 ± 0.025** | **-0.132 ± 0.033 \*\*\*/**<br>**-0.112 ± 0.027 \*\*\* (1rm)** | 0.107 ± 0.039 | 0.048 ± 0.047 | 0.479 ± 0.42 /<br>0.003 ± 0.067 (2rm) |

Results are reported twice; once including all outliers and once after removal of outliers 3sd from the overall mean. Different symbols are used for indicating statistical comparisons within the datasets that are significant (highlighted in bold): differences between controls and medicated patients in Dataset A and B use asterisks (\*\*, $p < .01$ ; \*\*\*, $p < .001$ ). There were no statistically significant differences between any controls and unmedicated patients in Dataset B nor within patients for their follow-ups (i.e. Dataset B unmedicated vs. medicated). Dataset A had all data of all controls and patients available (all $n=40$ ). Dataset B partly lacked finger tapping data: Controls, (baseline $n = 40$ , training $n = 40$ , consolidation, $n = 38$ ), Unmedicated (baseline $n = 37$ , training $n = 37$ , consolidation $n = 36$ ), Medicated (baseline $n = 36$ , training $n = 33$ , consolidation, $n = 35$ ). Further exclusion indicated with 1/2 rm = one/two outlier(s) more than 3sd from the mean removed.

### Sleep electroencephalography and subjective sleep quality

All patients and controls of all three datasets slept in the sleep laboratory of the Max Planck Institute of Psychiatry, Munich. All had an adaptation night before the study proper. Polysomnography was recorded (sampling rate of 200 or 250 Hz), stored and digitized (Dataset A and C: Comlab 32 Digital Sleep Lab, Brainlab V 3.3 Software, Schwarzer GmbH, Munich, Germany; Dataset B: 128 Ag/AgCl electrode setup [29], JE-209A amplifier, Neurofax Software, Nihon Kohden Europe GmbH, Rosbach, Germany) including EEG (Dataset A: filtered at 0.5–70 Hz, Dataset B: 0.016 Hz high pass only, Dataset C: filtered at 0.53-70 Hz), electrooculography (EOG), mental/submental electromyography (EMG) with a ground electrode attached at the forehead. Sleep was scored by independent experts (Dataset A: Rechtschaffen & Kales standards, Dataset B: AASM standards [30], Dataset C: Rechtschaffen & Kales standards using the Polysmith Software, Nihon Kohden Europe GmbH, Rosbach, Germany). Sleep stage 3/N3 and stage 4 were combined to SWS similar to the AASM standard and we use the latter for reporting, i.e. stage 1 as N1 and stage 2 as N2.

Patients and controls of Dataset B filled in the Pittsburgh Sleep Quality Index (PSQI) questionnaire as a self-report measure of sleep quality reporting (global score between 0 and 21, where a higher score reflects a worse subjective sleep quality).

### Depression severity

In Dataset A, depression severity was measured with self-rating instrument (BDI scores) for all patients as well as with a clinician rating instrument (Hamilton; HAMD scores) for a selection of 33 patients. In Dataset B and C, depression severity was measured with HAMD scores for all patients. For Dataset B, scores were measured at baseline (first EEG recording, unmedicated), after 7 days (second EEG recording, medicated) and after 28 days (medicated). For Dataset C, scores were measured at baseline, after 7 days (first EEG recording, medicated) and after 28 days (second EEG recording, medicated).

### Sleep EEG analysis

Analysis of sleep spindles and slow waves (SW) and SW-spindle coupling was performed using SpiSOP (https://www.spisop.org; RRID: SCR_015673), run in MATLAB 2013b (Mathworks, Natick, USA). All EEG analyses were performed on C3 and C4 leads (re-referenced to contralateral mastoids), as they were available in all three datasets. EEG signals were resampled at 100 Hz and low-pass filtered at 35 Hz. All parameters for power spectral analyses, sleep spindles, and SW were as reported in [31] and are described in brief in the following.

Power spectra were calculated using 5-s intervals that overlapped by 4 s for the Welch method resulting in frequencies of 0.6–30 Hz in a resolution of 0.2 Hz. To compare spectral power density between and within groups, for each 0.2 Hz frequency bin a permutation t-test was performed between each group pair with 10’000 simulations to account for multiple testing. Prior to statistical evaluation and for visualization, the power values were added 1 and dB-transformed.

Individual fast spindle center peak frequencies for detection were visually determined using power spectrum bands or when detection was unclear, the group average was used. Center peak frequencies were on average at 13.32 ± 0.08 (SE) Hz for Dataset A and 13.59 ± 0.07 Hz for Dataset B and 13.55 ± 0.08 Hz for Dataset C. Center peak frequencies did not differ for any two-group comparisons in each dataset. We focused the analysis on fast sleep spindles only, as power peaks of slow spindles could not be clearly identified in too many participants for the central channels (most recordings lacked frontal channels) and less reliability of detection especially in older individuals lacking SWS. Fast spindles are denoted as spindles throughout the paper for brevity.

The SW detection targeted a frequency range of 0.5–1.11 Hz with resulting core frequencies of ∼0.75 Hz as the major contributor to SWS and SWA (i.e. the non-REM-typical slow waves of larger amplitudes). We report both spindle and SW densities and counts, amplitudes, and duration.

Parameters for coupled sleep spindles to SW (SW-spindles) were as described elsewhere [32], with the exception that SW was identified with a factor of 1.25 for the means of the amplitude and the negative half-wave peak potential, and only one threshold of 1.5 standard deviations of the filtered signal to mark spindles; the SW-spindles were identified only with spindles falling in a time window from the peak down-state to the end of the SW (i.e. the next up-to-down zero crossing) to better target (fast) sleep spindles. Sleep spindles were counted only once for the first slow-wave in which they occurred within the same channel. The mean delay of sleep spindles to the SW down-state and the standard deviation of this delay were calculated to estimate the temporal dispersion of their co-occurrence (delay dispersion). In addition, the average amplitude and duration of coupled SW and spindles were calculated. We chose these measures, as they capture the potential variation of SW-spindle coupling sufficiently for our aims. In addition, these measures do not assume sinusoidal progression of oscillations. This is particularly important as measures that rely on the phase angle of oscillations (like phase-locking values or modulation index) are heavily confounded by overly simplistic assumptions that those brain waves are continuous, homogenous waves and are thus harder to interpret [33]. Epochs with EMG and EEG artifacts and channels with more than 20% artifacts during non-REM sleep were manually excluded by an experienced scorer before all automatic analyses.

All analyses files including R-scripts and SpiSOP files are made public and can be accessed under https://osf.io/bdez9/.

### Statistical Analysis

All statistical analyses were performed using the R programming language (version 3.5.1; R Core Team, 2018) and MATLAB 2015b (Mathworks, Natick, USA). The differences between MDD patients and controls were analyzed with two-tailed independent t-tests, whereas the differences between MDD patients within the datasets were analyzed with dependent t-tests. Variances were assumed unequal unless otherwise specified. Outliers that were 3 standard deviations from the mean were automatically removed for the separate analyses. Details on outlier removal per analysis are added in the result section. All statistical analyses were performed using functions from base R or using the R-package “*rstatix”* [34].

Bayes factors (BF) were calculated using the R-package *“BayesFactor”* [35] for all statistical tests as a statistical quantification of the evidence for the alternative over the null hypothesis. A BF_10_ less or equal to 1 quantifies relative evidence in favor of the null hypothesis (H0), while a BF_10_ larger than 1 quantifies relative evidence for the alternative hypothesis (H1). BF_10_ values could be interpreted as either anecdotal (1–3), moderate (3–10), strong (10–30), very strong (30–100), or extreme (> 100) evidence for H1. All BFs were calculated with default uniform prior scales (r scale = 0.5). The first draft of this paper was created using R markdown (R Core Team, 2018) and the R-package *“papaja”* [36]. All plots were created using the R-package “*ggplot2*” [37] and inspired by the RainCloud plot [38]. Mean values are given ± their standard error. Please note that we did not correct the statistical tests for multiple comparisons as the study aimed to be explorative and descriptive in nature, nevertheless we adopted corrective measures on a per analysis basis if indicated (i.e. permutation testing).

## Results

### Sleep architecture and quality

For differences in percentages of all datasets see Table 2. For all statistical comparisons and details, see supplemental Table S1.

**Table 2.** Sleep architecture table (mean ± SE)

|  | Dataset A |  | Dataset B |  |  | Dataset C |  |  |
| --- | --- | --- | --- | --- | --- | --- | --- | --- |
|  | Controls | Medicated | Controls | Unmedicated | Medicated 7d | Controls | Medicated 7d | Medicated 28d |
| N1[%] | **8.45 ± 0.75** | **13.9 ± 1.25<sup>\*\*\*</sup>** | **13.8 ± 0.95** | **14 ± 1.08** | **17.4 ± 1.08<sup>\*/++</sup>** | **9.89 ± 0.62** | **14 ± 1.21<sup>\*\*</sup>** | **14.7 ± 1.42<sup>\*</sup>** |
| N2 [ %] | 46.2 ± 1.43 | 50.3 ± 1.83 | 45.4 ± 1.11 | **43.5 ± 1.23** | **47.1 ± 1.47<sup>++</sup>** | 48.7 ± 1.58 | 49.7 ± 2.18 | 49 ± 1.69 |
| SWS [%] | 16.2 ± 1.33 | 13.1 ± 1.64 | 19.1 ± 1.18 | 17.6 ± 1.31 | 16.1 ± 1.26 | 15.4 ± 1.41 | 13.4 ± 1.7 | 13.9 ± 1.8 |
| Non-REM [ %] | 62.3 ± 1.26 | 63.4 ± 1.86 | 64.5 ± 1.06 | 61.1 ± 1.59 | 63.2 ± 1.66 | 64.2 ± 1.32 | 63.1 ± 1.63 | 62.9 ± 1.72 |
| REM [ %] | **18.6 ± 0.83** | **14.3 ± 1.38<sup>\*\*</sup>** | **17 ± 0.915** | **16.8 ± 0.92** | **12.1 ± 1.01<sup>\*\*\*/+++</sup>** | **18.3 ± 0.85** | **13 ± 1.43<sup>\*\*</sup>** | **14 ± 1.45<sup>\*</sup>** |
| WASO [ %] | 10.4 ± 1.2 | 7.8 ± 0.9 | **4.66 ± 0.53** | **8.16 ± 1.16<sup>##</sup>** | **7.33 ± 0.96<sup>\*</sup>** | 7.27 ± 1.2 | 9.09 ± 1.13 | 7.28 ± 0.98 |
| TST [min] | 461 ± 4.29 | 462 ± 3.6 | 471 ± 2.11 | 468 ± 4.56 | 468 ± 2.41 | 467 ± 3.67 | 464 ± 2.81 | 466 ± 2.44 |
| Sleep onset [min] | 22.7 ± 2.68 | 21.7 ± 2.12 | 15.7 ± 1.55 | 24.1 ± 5 | 20.9 ± 2.24 | **26.3 ± 1.4** | **32.4 ± 2.57<sup>\*</sup>** | 30.9 ± 2.61 |
| SWS onset [min] | 21.5 ± 3 | 31.6 ± 4.84 | 18.4 ± 1.42 | 26.8 ± 4.15 | 26 ± 3.67 | 19 ± 2.35 | 24.1 ± 3.54 | 19.7 ± 2.12 |
| REM onset [min] | **74 ± 5.08** | **177 ± 17.5<sup>\*\*\*</sup>** | **89.4 ± 5.76** | **105 ± 9.18** | **174 ± 14.7<sup>\*\*\*/+++</sup>** | **79.2 ± 5.91** | **186 ± 19.9<sup>\*\*\*</sup>** | **173 ± 18.8<sup>\*\*\*</sup>** |

Sleep stage percentages are given with respect to total sleep time (TST). Note that non-REM sleep was defined as the combination of N2 and slow-wave sleep (SWS) without N1 sleep. Different symbols are used for indicating statistical comparisons within the datasets that are significant (highlighted in bold): differences between Controls and Medicated patients in Dataset A, B, and C use asterisks (\*, $p < .05$ ; \*\*, $p < .01$ ; \*\*\*, $p < .001$ ), between Controls and Unmedicated patients in Dataset B, use hashes (##, $p < .01$ ), and within patients for their follow-ups ( Dataset B unmedicated vs. medicated 7d) use pluses (++, $p < .01$ ; +++, $p < .001$ ).

In Dataset A, compared to controls, the medicated MDD patients had a higher proportion of N1 sleep, (*t*(63.93) = 3.71, *p* <.001), but a lower proportion of REM sleep (*t*(63.70) = -2.71, *p* = .009) and took longer time to reach it (*t*(45.48) = 5.65, *p* <.001). They also had a slightly higher proportion of N2 sleep that did not reach significance (*p* = .082), as well as a longer SWS latency, which also did not reach statistical significance (*p* = .08). The increase in N1 did not account well for the loss of REM sleep in MDD patients (*r* = -.26, *p* = .102). The total sleep time (TST) did not differ between patients and medicated MDD patients.

In Dataset B, compared to controls, the unmedicated MDD patients spend a higher proportion awake after sleep onset (*t*(51.64) = 2.74, *p* = .008). Patients spent less time in non-REM sleep, and SWS onset occurred later, but both did not reach significance (*p* = .08, *p* = .06 resp.). There were no other group differences in the proportion spent in any sleep stage between unmedicated patients and controls. Also under medication a week later, MDD patients, compared to controls, spent a higher proportion awake after sleep onset (*t*(57.12) = 2.41, *p* = .019) and also a higher proportion in N1 sleep (*t*(74.13) = 2.49, *p* = .015). In addition, they spent a lower proportion of sleep in REM (*t*(74.88) = -3.63, *p* = .001) and took longer to reach it (*t*(48.16) = 5.38, *p* <.001). Less time was spent in SWS, and patients took longer to reach it, but both did were not statistically significant (*p* = .09, *p* = .06 resp.). Similar to the differences between control participants and unmedicated patients, the patients during medication follow-up spent a higher proportion in N1 sleep (*t*(37) = 2.8, *p* = .008), as well as in N2 sleep (*t*(37) = 2.83, *p* = .008) than an unmedicated state a week prior. They also spent a higher proportion in REM sleep as well (*t*(37) = -4.39, *p* <.001) and took longer to reach it (*t*(37) = 4.3, *p* < .001). The total sleep time (TST) did not differ within patient groups or to controls. Here, in the medicated patients, the increase in N1 was positively related to the loss of REM sleep (*r* = -.34, *p* = .04).

Controls (*n* = 38) rated their subjective self-assessed sleep quality (PSQI: 3.97 ± 0.3 (SE)) markedly different from unmedicated (*n* = 32, 10.2 ± 0.68, *t*(43.1) = -8.46, *p* < .001, BF_10_ > 100), or medicated patients (*n* = 35, 8.74 ± 0.643, *t*(48.34) = -6.72, *p* < .001, BF_10_ > 100). Patients rated their sleep of similar quality in unmedicated as in medicated state (*p* = .12). Across groups, the PSQI scores were weakly associated with sleep onset (*r* = .2, *t*(103) = 2.08, *p* = .04).

In Dataset C, compared to controls, the 7-day medicated patients spent a higher proportion in N1 sleep (*t*(42.89) = 3.03, *p* = .004), less proportion in REM sleep (*t*(46.82) = -3.16, *p* = .003), and took longer to reach it (*t*(34.03) = 5.13, *p* < .001). They also took longer to reach sleep onset (*t*(44.64) = 2.09, *p* = .042). Similarly, compared to controls, the 28-day medicated patients spent a higher proportion in N1 sleep (*t*(37.45) = 3.11, *p* = .003), less proportion in REM sleep (*t*(46.82) = -3.16, *p* = .015) and took longer to reach it (*t*(34.03) = 5.13, *p* < .001). Notably, no group differences were found between the 7-day medicated patients and the same patients after 28 days. Similarly to Dataset B, in the 7-day as well as the 28-day medicated patients, the increase in N1 was positively related to the loss of REM sleep (*r* = -.56, *p* = .001, *r* = -.44, *p* = .02, resp.).

In summary, in each dataset, all medicated patients had a higher proportion of light sleep in N1 and a lower proportion of REM sleep and longer REM latency, which was not observed in the only available unmedicated state (Dataset B). Supplementary sleep cycle analysis (equivalent to [39]) confirmed these findings and revealed that medicated patients across datasets showed extended 1^st^ and 2^nd^ sleep cycles resulting from longer non-REM periods. In addition, the long-term medication groups (i.e. Dataset A Medication, Dataset C Medicated 28d) had prolonged REM periods in these early cycles which seemed to be driven by patients receiving REM suppressing medication (see supplemental Table S4). Unmedicated patients (Dataset B) showed similar results compared to controls except for increased wake after sleep onset, which seemed increased independent of medication state and was also not observed in the other datasets. Importantly, no group differences in the sleep architecture for N2, SWS, or combined non-REM sleep was found for each dataset, except for the patients in Dataset B in unmedicated vs medicated state. Combining the three (medicated and control) datasets, however, revealed a reduction in SWS proportion as well as an increased SWS latency in the medicated vs control subjects (see supplementary materials for details).

### Spectral power

In Dataset A, medicated MDD patients showed reduced power in the lower frequency ranges (1–4.6 Hz) overlapping with the SWS range (0.5–4 Hz) of non-REM sleep (Figure 2A).

**Figure 2:**
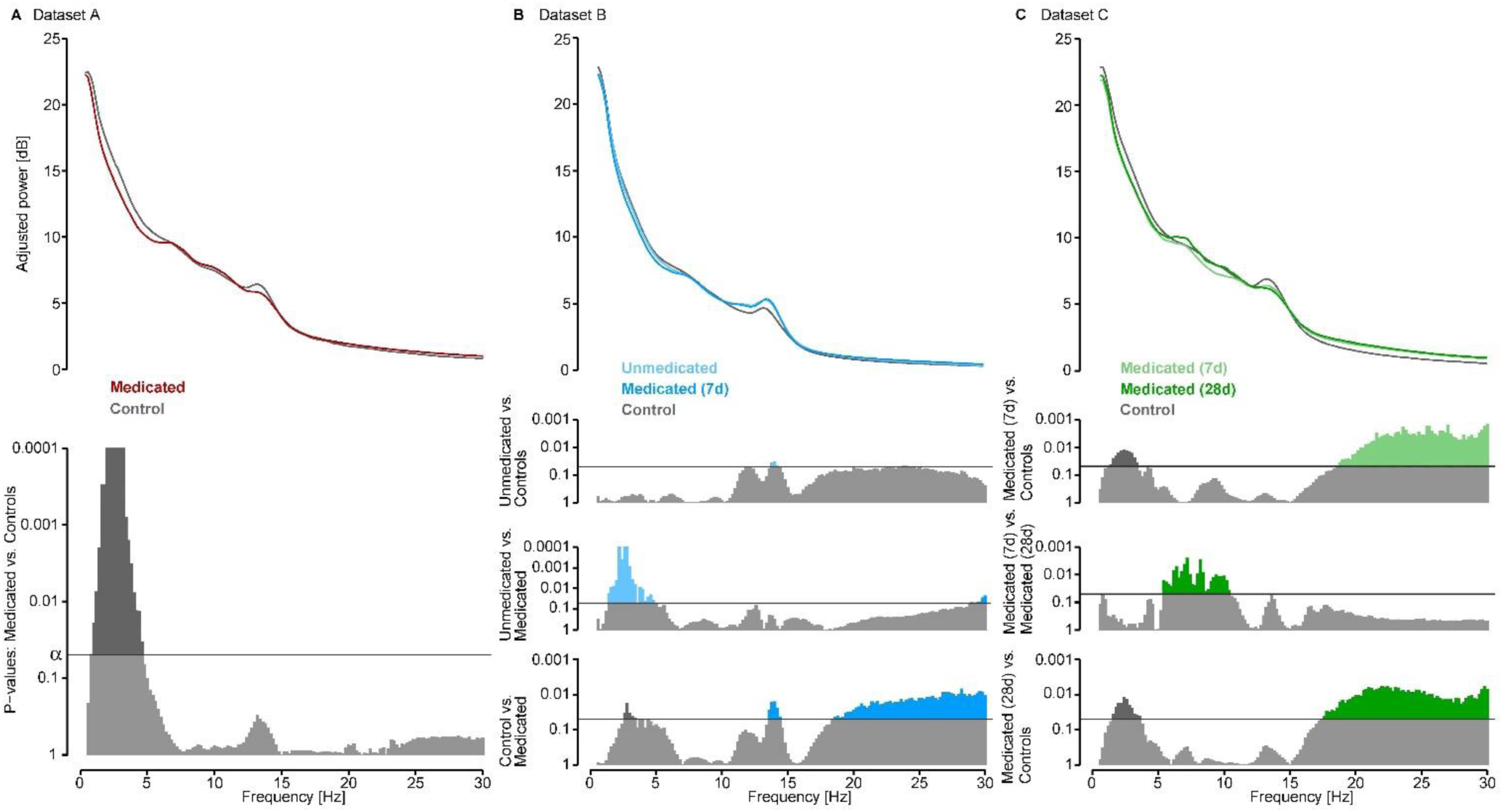
EEG non-REM sleep power spectra group comparisons. (**A**) In Dataset A, Medicated MDD patients (red) show reduced power in 1–4.6 Hz of non-REM sleep compared to controls (grey). (**B**) In Dataset B, Unmedicated MDD patients compared to controls (grey) show increased power in the fast spindle band range (13.6–14.4 Hz). Unmedicated MDD patients compared to themselves in the medicated state (dark blue) show increased power in the lower frequencies (2.6–3.4Hz). Medicated patients, compared to controls show reduced power in lower frequencies but increased power in the fast spindle band as well as in the higher frequencies (>18 Hz). (**C**) In Dataset C, both 7-day (light green) and 28-day (dark green) medicated MDD patients show reduced power in the lower frequencies (1.2–3.4 Hz and 1.6 –3.6 Hz resp.) and increased power in the higher frequencies (>18.6 Hz and > 17.4 Hz resp.) compared to controls (grey). In addition, 7-day medicated MDD patients show reduced power in the 5.5–10.4 Hz range compared to the same patients after 28 days.

In Dataset B, medicated MDD patients had reduced non-REM power in the lower frequencies (2.6–3.4 Hz) but also higher power in the fast spindle band (13.6–14.6 Hz) and >18.6 Hz frequencies compared to controls. In contrast, unmedicated MDD patients had reduced power mainly in the spindle frequency range (13.6–14.4 Hz) and increases in higher frequencies (around 20 and up to 27 Hz) compared to controls (Figure 2B).

In Dataset C, 7-day medicated patients showed lower non-REM power in lower frequencies (1.2–3.4 Hz), as well as in higher frequencies (>18.6 Hz) and similarly between controls and 28-day medicated patients (1.6–3.6 Hz and > 17.4 Hz). Patients showed a decrease in the 7-day medicated patients in higher theta and alpha bands (5.5–10.4Hz) power in non-REM sleep compared to their 28-day medicated follow up (Figure 2C).

In summary across datasets, medicated MDD patients had lower power in the higher SWA bands, which was not the case for the unmedicated patients (Dataset B). There were power increases in the spindle band in dataset B for both the medicated and unmedicated patients that was not observed in the other datasets. Spurious reductions in alpha activity and alterations in higher frequency bands were observed but not consistent within and across dataset

### Non-REM sleep events

We characterized occurrences and properties of hallmark non-REM sleep events, i.e. (fast) sleep spindles, slow waves (SWs), and their coupling in SW-spindles. For an overview of all reported values all datasets see supplemental Table S2 and for all statistical comparisons and details, see supplemental Table S3.

#### Sleep spindles

In Dataset A, only spindle density (per epoch) in MDD patients was slightly, but not significantly, decreased compared to controls (*p* = .052) reaching significance after removal of 1 outlier 3 SD below the mean in the control group (*t*(70.79) = -2.58, *p* = .012). No group differences in spindle count, amplitude, or duration were found (Figure 3A).

**Figure 3:**
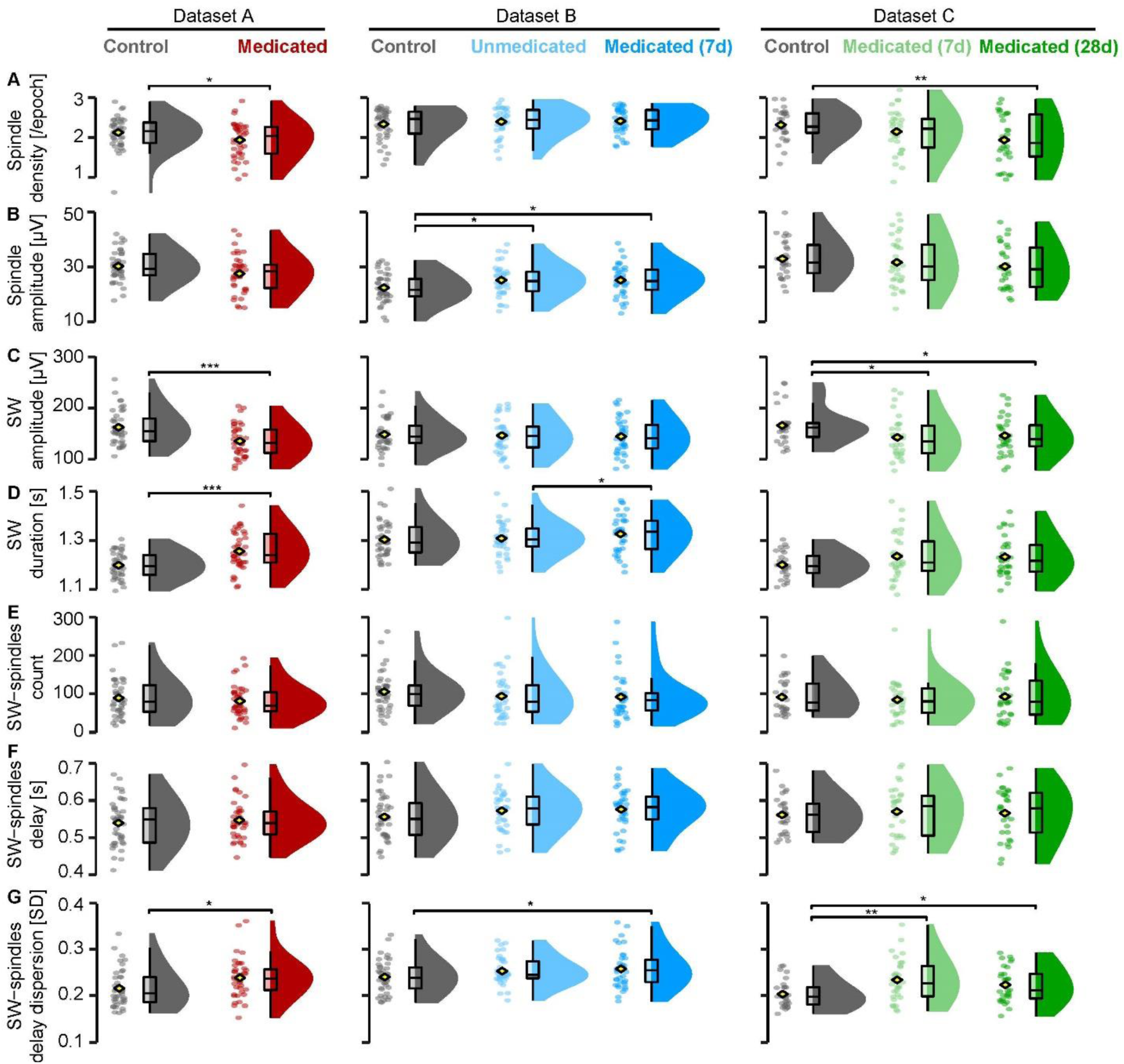
Non-REM event features. (**A**) In Dataset A, Medicated MDD patients had a lower sleep spindle density than Controls (outlier depicted, significance based after outlier removal), as do 28-day Medicated patients compared to Controls in Dataset C. (**B**) In Dataset B, Unmedicated MDD patients show higher sleep spindle amplitudes than Controls. (**C**) In Dataset A, Medicated MDD patients show lower slow waves (SW) amplitudes than Controls, as did 7-day and 28-day medicated patients compared to controls in Dataset C. (**D**) In Dataset A, Medicated MDD patients had SW of longer duration than Controls, as did Medicated compared to Unmedicated patients in Dataset B. (**E**). No differences in each dataset on SW-spindle counts (**F**) nor delay between spindle and SW (downstate) trough (**G**) but an overall increase in delay dispersion (spread of the delay, in standard deviation [SD]) of spindles within SW in Medicated compared to Controls can be seen across datasets. Data is depicted in clouds of dots per individual, group mean (yellow diamond), and a 25%–75% quarter-quartile boxplots with minimum and maximum the 1.5 times the inter-quartile difference and smoothed density plots within the full group range. Significances for two-group comparisons in asterisks (*, *p* < .05; **, *p* < .01, ***, *p* < .001).

In Dataset B, patients showed a higher spindle amplitude than controls in unmedicated (*t*(75.06) = 2.21, *p* = .03) and medicated state (*t*(75.12) = 2.22, *p* = .03). In addition, medicated patients show a longer spindle duration, but this did not reach significance (*p* = .051). No other group differences in spindle parameters were observed (Figure 3B).

In Dataset C, spindle density (per epoch) in 28-day medicated MDD patients was decreased (*t*(49.21) = -2.79, *p* = .007; Figure 3A), as well as spindle count (*t*(51.94) = -2.5, *p* = .016) and spindle duration (*t*(55.98) = -2.44, *p* = .018) compared to controls.

In summary across datasets, spindle density was reduced in the MDD patient groups with the longest time of medication (Dataset A and Dataset C 28-day medication follow-up) but not after medication of one week (Dataset C 7-day and Dataset B medicated) compared to controls.

Consistent with the spindle band power increases, the spindle amplitude was increased for the unmedicated sample in Dataset B, but did not change from those levels with the short-term medication follow-up. No differences in spindle parameters were found when all three (medicated and control) datasets were combined (see supplementary materials for details).

#### Slow waves

In Dataset A, the medicated MDD patients showed overall a lower SW amplitude than healthy controls (Figure 3C, *t*(77.69) = -3.75, *p* < .001), a longer SW duration (Figure 3D, *t*(68.6) = -3.86, *p* < .001) and consequently a lower SW frequency (*t*(72.07) = -3.77, *p* < .001). No group differences in SW density or count were found.

In Dataset B, no group differences in SW density, count, amplitude were found between any groups. Only the medicated compared to the unmedicated patients showed a longer SW duration (Figure 3D, *t*(37) = 2.32, *p* = .026) and a lower SW frequency (*t*(37) = -2.33, *p* = .025).

In Dataset C, compared to controls both 7-day medicated as well as 28-day medicated patients showed decreased SW amplitude, respectively (Figure 3C, *t*(55.79) = -2.33, *p* = .024; *t*(55.95) = -2.07, *p* = .043). SWs were also generally longer in MDD patients compared to controls (Figure 3D), but did not reach statistical significance (*p* > 0.09 [vs 7d], *p* > 0.07 [vs 28d]).

In summary across datasets, consistent with the observed reduced power in the SWA band and lower amounts of SWS, SW amplitudes were reduced in medicated MDD patients. However, although this effect was present even in short-term medication (1 week) of Dataset C, this was not observed in patients with a younger age distribution (Dataset B). SW duration was longer in the youngest medicated sample compared to their unmedicated state (Dataset B) and increased under medication compared to controls (Dataset A). When all (medicated and control) datasets were combined, MDD patients showed longer and smaller (reduced amplitude) SWs (see supplementary materials for details).

#### SW-spindle coupling

Because SW-spindle coupling has repeatedly been associated with memory consolidation and spectral power analysis indicated patient-control group differences in both the SWA and spindle range, we checked for group differences in SW-spindle coupling. Please note, the longer duration in SW might increase the chances for spindles to align (Figure 3D). No group differences in the SW-spindle counts nor the delay were found in any of the datasets (Figure 3E, F).

In Dataset A, MDD patients showed a greater delay dispersion – or spread around the delay (*t*(77.89) = 2.46, *p* = .016). In Dataset B, medicated MDD patients showed a greater delay dispersion than controls (*t*(73.76) = 2.02, *p* = .047). In Dataset C, both 7-day medicated MDD patients as well as 28-day medicated MDD patients showed a greater delay dispersion than controls (Figure 3G, *t*(49.61) = 3.03, *p* = .004; *t*(54.5) = 2.28, *p* = .03, resp.).

Furthermore, we calculated the properties of *coupled* and *uncoupled* spindles (amplitude, duration, frequency) and SW (amplitude, duration) and explored if they differed per group in all datasets. In Dataset A, MDD patients, compared to controls, showed lower amplitude (*t*(76.47) = -3.22, *p* = .002) as well as longer duration of SW that coupled with a spindles (*t*(71.75) = 3.37, *p* = .001). In Dataset B, medicated MDD patients, compared to controls, showed higher spindle amplitude of those spindles that coupled with a SW (*t*(75.32) = 2.27, *p* = .026). In addition, medicated patients compared to controls, showed a larger difference in spindle frequency (*t*(68.99) = -2.39, *p* = .02) as well as spindle duration (*t*(75.95) = -2.1, *p* = .039) between coupled and uncoupled spindles. In Dataset C, 7-day medicated MDD patients, compared to controls, showed longer spindle duration of those spindles that coupled with a SW, (*t*(54.77) = 2.53, *p* = .014), as well as lower amplitude and longer duration of SW that coupled with a spindles, (*t*(55.88) = -2.63, *p* = .011; *t*(50.29) = 2.8, *p* = .007, resp.). Patients after 28 days of medication compared to 7 days of medication showed a shorter spindle duration in the coupled spindles, (*t*(57.53) = 2.05, *p* = .045). Similarly, 28-day medicated MDD patients, compared to controls, lower amplitude and longer duration of SW that coupled with a spindles, (*t*(55.73) = -2.3, *p* = .025; *t*(52.54) = 2.56, *p* = .013, resp.). 7-day medicated MDD patients, compared to controls, showed a smaller difference in spindle duration between coupled and uncoupled spindles, (*t*(55.31) = 2.72, *p* = .009). In addition, 7-day medicated patients compared to 28-day medicated patients showed a larger difference in SW duration between coupled and uncoupled SW (*t*(55.84) = 2.12, *p* = .039). Similarly, 28-day medicated MDD patients, compared to controls, showed also a smaller difference in spindle duration between coupled and uncoupled spindles, (*t*(55.24) = - 2.43, *p* = .02). For an overview of all reported values all datasets see supplemental Table S2 and for all statistical comparisons and details, see supplemental Table S3. Lastly, we combined the three datasets by pooling all controls (*n* = 108) and all the patients in a medicated state (*n* = 108). All the above-reported analyses were repeated on the combined data. See supplementary materials and Table S5 for more details.

In summary across datasets, not indicated by the observed spindle or SW features alone, there was a consistent increase in delay dispersion of spindles within SW in medicated MDD patients against controls that also went along with prolonged spindles in SW and – depending on the dataset – decreased amplitude and increased duration of spindle-coupled SW.

### Medication

In all datasets, when medicated, the MDD patients were prescribed a great variety of antidepressant medication classes and most took a combination of different types. In addition, some took benzodiazepines and GABA-ergic drugs (typically hypnotics), which are known to influence sleep [40]. Given the variety of drugs, no analysis on the specific type of medication on any of the outcome measurements of interest were performed on the specific datasets. However, medication data was available for 103 patients after combining the three datasets (all the patients in medicated state were pooled together, for Dataset C the patients after 7 days of medication were taken) that could be pooled into 5 subgroups by their main medication type: selective serotonin reuptake inhibitors (SSRIs, n = 32), serotonin-norepinephrine reuptake inhibitors (SNRIs, n = 34), tricyclic antidepressants (TCAs, n = 33), hypnotics (n = 11), or an alternative drugs (n = 47). We then compared those subgroups against their pooled controls on representative sleep parameters (i.e. spindle density, spindle count, SW amplitude, SW duration, non-REM percent, REM percent, SW-spindle count, SW-spindle delay, SW-spindle delay dispersion and procedural memory consolidation).

Spindle count was descriptively higher in patients taking hypnotics, but this did not reach significance (*p* =.1) and SW amplitude was decreased (*b* = -25.05, *t*(101) = -2.35, *p* = .021). Since hypnotics were only prescribed in Dataset A and C, which included older patients, we added age as a moderating predictor in the regression model as age is known to decrease SW amplitude [41]. The interaction effect between the hypnotic and age on SW amplitude was not significant, suggesting age was not a strong mediator of the SW amplitude decreases on hypnotics (*p* = 0.12). As expected SW amplitude was reduced with age (main effect, *b* = -1.37, *t*(99) = -6.38, *p* < .001). Lastly, TCAs prescription was associated with a decrease in SW amplitude as well, (*b* = -16.46, *t*(101) = -2.33, *p* = .022) and age was no moderator on this association (*p* = 0.9).

In addition, we explored the influence of the three most common prescribed specific medication drugs in our combined sample. These were venlafaxine (SSNRI, *n* = 24), mirtazapine (NaSSA, *n*= 19) and trimipramine (TCA, *n* = 18). Of these drugs, only venlafaxine showed significant influences, namely decreased spindle density (*b* = -0.23, *t*(108) = -2.08, *p* = .039) as well as, expectedly, decreased the time spent in REM sleep (*b* = -6.82, *t*(108) = -4.16, *p* < .001).

### Procedural memory

Finger tapping data was only available in Dataset A and Dataset B. Here, medicated MDD patients (as previously published in 28) initially tapped less sequences correctly on the first baseline run than controls in Dataset A (ΔM = 2.50, 95% CI [1.03, 3.97], *t*(74.60) = 3.38, *p* = .001, BF_10_ = 27.41, one outlier removed: *t*(76.57) = 3.23, *p* = .002). No such baseline differences were observed in Dataset B for the unmedicated vs control (*p* = .2, BF_10_ = 0.46) nor the medicated vs control (*p* = .8, BF_10_ = 0.25). Both datasets A and B, did not differ in their training performance, or learning effect (i.e. difference the first and the mean of the final three runs), *p* >.2. Importantly, overnight consolidation (i.e. difference between the mean of the final three runs before sleep and the three runs after sleep) in medicated MDD patients was impaired compared to controls in Dataset A (ΔM = 0.22, 95% CI [0.14, 0.30], *t*(72.36) = 5.26, *p* < .001, BF_10_ >100, one outlier removed: *t*(76.61) = 5.41, *p* < .001) but not in the unmedicated vs control group in Dataset B (*p* = .5, BF_10_ = 0.31), nor for the medicated vs control group of Dataset B (*p* = .9, BF_10_ = 0.25. two outliers removed; *p* = .2, BF_10_ = 0.42). Between the unmedicated and medicated state of the same MDD patients, there were no group differences at baseline, in training effect, nor in overnight consolidation (all *p* > 0.1, all BF_10_ < 0.55). See Figure 4 and Table 3 for all group differences. Again, we repeated the analysis on the combined behavioral data of Dataset A and B. See supplementary materials, Table S6 and Figure S2 for more details.

**Figure 4:**
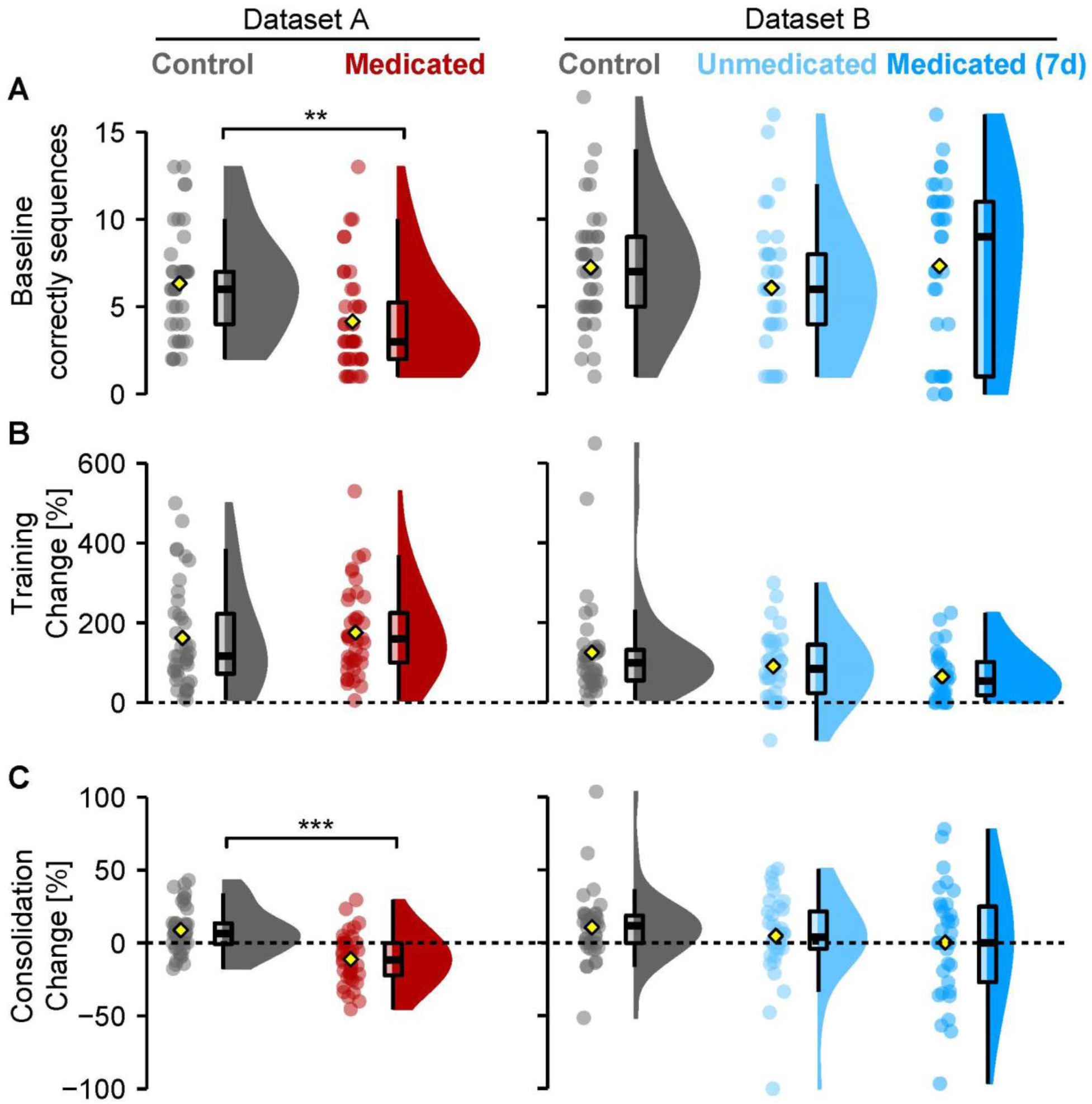
Finger tapping task performance for baseline, training, and consolidation over 24 hours. (**A**). Amount of correctly tapped sequences of the first 30-second run. Medicated patients in Dataset A tap fewer sequences correct than Controls. (**B**) Percentage change score between the first run and the mean of the last three runs. (**C**) Percentage change score between the mean of three test runs after sleep in the morning and the mean of the last three training runs before sleep. Medicated patients in Dataset A perform worse after sleep than Controls. Data depicted like in Figure 3. Significances for two-group comparisons in asterisks (**, p < .01; ***, p < .001).

In summary, procedural memory impairments at baseline and consolidation in mainly long-term medicated older MDD patients observed previously in Dataset A were not observed in the younger age group of patients in Dataset B for either unmedicated or short-term medicated state.

### Sleep parameters related to overnight consolidation performance

As procedural memory data was only available in Dataset A and Dataset B, the following part will exclude Dataset C. No interaction effects between group (control and patient) and sleep architecture parameters were found.

#### Spectral power

Given the significant group differences in the delta range in Dataset A, we checked if this group difference was related to overnight consolidation performance. However, no interaction effect between group and average SWA/delta power between 1–4.6Hz on overnight consolidation was significant, nor in Dataset B (*p* > 0.9). In addition, no between-group (controls and medicated state) interaction with average sigma power between 13.6-14.6Hz on overnight consolidation (*p* > 0.7) was found.

#### Sleep Spindles

In Dataset A, given the already lowered spindle density in patients, their impaired consolidation for finger tapping compared to controls seemed to be less pronounced with increasing spindle density (*b* = 0.19, *t*(76) = 2.07, *p* = .042). After removing the same outlier in the patient group as before, however, this interaction effect vanished (*p* = .21). No other interaction with other spindle parameters (count, amplitude, duration, frequency) reached significance.

In Dataset B, no interactions between spindle density, count, amplitude, duration or frequency, and group (between controls, unmedicated and medicated) on overnight consolidation were found.

#### SW-spindle coupling

Lastly, we checked if any of the group differences in the coupling parameters were correlated with overnight consolidation. In Dataset A, an interaction between SW-spindle count and group on overnight consolidation (after removal of one outlier) was found (*b* = 0.002, *t*(75) = 2.02, *p* = .047), which indicated a stronger positive correlation between the SW-spindle count and overnight consolidation for patients (*r* = 0.422), than for controls (*r* = 0.04). Similarly, an interaction between SW-spindle delay dispersion and group on overnight consolidation (after removal of the same outlier) was found, (*b* = -2.27, *t*(75) = -2.47, *p* = .015). Here, patients with more delay dispersion (i.e., more temporal variance between the spindle and SW co-occurring) showed worse overnight consolidation (*r* = -0.295), whereas this relation was less strong in controls (*r* = 0.25).

In Dataset B, no significant interaction effect between SW-spindles count, delay its dispersion, and group between any of the groups (*p* > 0.4).

### Depression severity and outcome

Depression severity (HAMD) for patients in each of the three datasets, is shown in Table 4. In both datasets B and C, depression scores significantly decreased between baseline and 7 days, (*p* < .001), as well as between 7 days and 28 days, (*p* < .001). We combined the three datasets to check for a relationship between depression severity and spindle density, count, slow-wave amplitude, duration, SW-spindle delay dispersion, and overnight. All the controls were pooled together, as were all the patients in a medicated state, which for Dataset C were the patients after 7 days of medication. HAMD scores administered at the time of the pooled EEG recording sessions were used in the analysis.

**Table 4.**
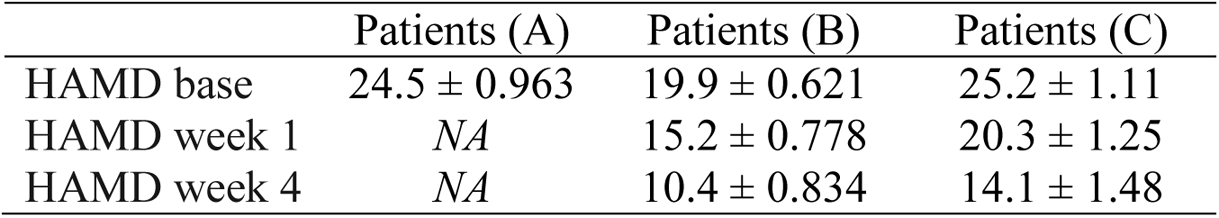
Depression severity over time in HAMD scores (mean ± SE)

Higher HAMD scores were related to less spindle density (*b* = -0.02, *t*(99) = -3.28, *p* = .001, BF_10_ = 21.51), lower spindle count (*b* = -12.98, *t*(99) = -2.46, *p* = .016, BF_10_ = 2.91) and shorter spindle duration (*b* = -0.002, *t*(99) = -2.98, *p* = .004, BF_10_ = 9.81). No association between the SWs or SW-spindle features and depression severity reached significance. In addition, no association between depression severity and overnight procedural memory consolidation was found (*p* > 0.1). To check if these links are explained by age, a moderation analysis was performed by adding age as a predictor in the regression model. The interaction between depression severity and age on spindle density was not significant (*p* > 0.6) suggesting age is not a strong moderator to the depression-severity-spindle effects. As expected, a main effect of age indicated that older patients show lower spindle density (*b* = -0.02, *t*(97) = -2.24, *p* = .03). In addition, we checked for similar associations between sleep parameters and the number of patient-reported total depression episodes. The number of episodes was negatively related to spindle frequency (*b* = -0.07, *t*(97) = -2.54, *p* = .012), as well as to SW amplitude (*b* = -3.6, *t*(97) = -2.36, *p* = .21). No association between number of episodes and overnight consolidation was found (*p* > 0.4). Lastly, Dataset B and C reported a binary classification of treatment response based on ≥ 50% reduction in Hamilton score between baseline and after 28 days (*n* = 68). None of the reported sleep features were significant predictors of treatment response. See Table 5 for a complete overview of all correlations between sleep parameters and depression outcome scores.

**Table 5.**
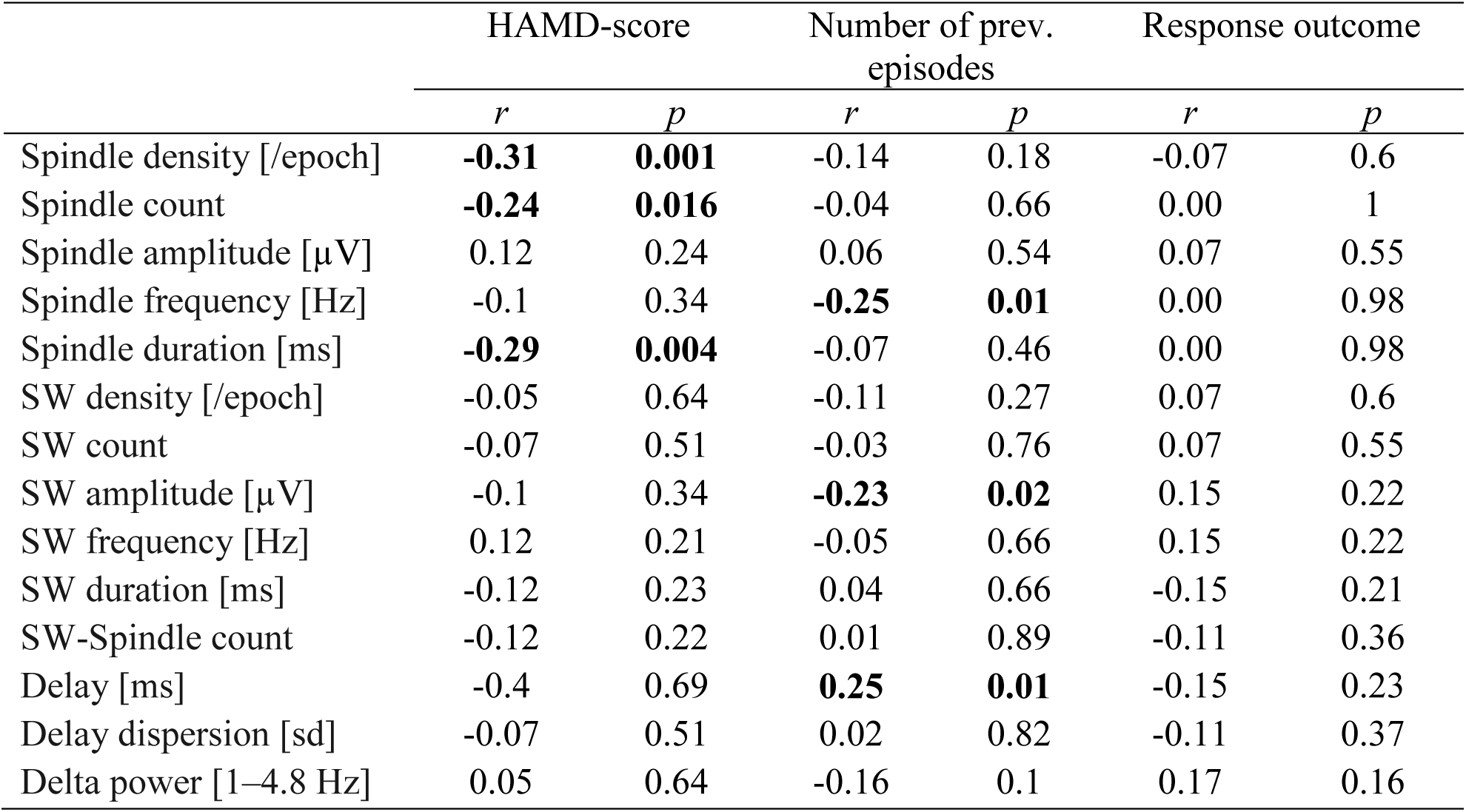
Correlation table between depressive scores and sleep parameters.

|  | HAMD-score |  | Number of prev. episodes |  | Response outcome |  |
| --- | --- | --- | --- | --- | --- | --- |
|  | *r* | *p* | *r* | *p* | *r* | *p* |
| Spindle density [/epoch] | **-0.31** | **0.001** | -0.14 | 0.18 | -0.07 | 0.6 |
| Spindle count | **-0.24** | **0.016** | -0.04 | 0.66 | 0.00 | 1 |
| Spindle amplitude [μV] | 0.12 | 0.24 | 0.06 | 0.54 | 0.07 | 0.55 |
| Spindle frequency [Hz] | -0.1 | 0.34 | **-0.25** | **0.01** | 0.00 | 0.98 |
| Spindle duration [ms] | **-0.29** | **0.004** | -0.07 | 0.46 | 0.00 | 0.98 |
| SW density [/epoch] | -0.05 | 0.64 | -0.11 | 0.27 | 0.07 | 0.6 |
| SW count | -0.07 | 0.51 | -0.03 | 0.76 | 0.07 | 0.55 |
| SW amplitude [μV] | -0.1 | 0.34 | **-0.23** | **0.02** | 0.15 | 0.22 |
| SW frequency [Hz] | 0.12 | 0.21 | -0.05 | 0.66 | 0.15 | 0.22 |
| SW duration [ms] | -0.12 | 0.23 | 0.04 | 0.66 | -0.15 | 0.21 |
| SW-Spindle count | -0.12 | 0.22 | 0.01 | 0.89 | -0.11 | 0.36 |
| Delay [ms] | -0.4 | 0.69 | **0.25** | **0.01** | -0.15 | 0.23 |
| Delay dispersion [sd] | -0.07 | 0.51 | 0.02 | 0.82 | -0.11 | 0.37 |
| Delta power [1–4.8 Hz] | 0.05 | 0.64 | -0.16 | 0.1 | 0.17 | 0.16 |

Lastly, given our other finding that unmedicated patients showed a high spindle amplitude compared to healthy controls, we explored if spindle amplitude was related to depression severity and outcome in unmedicated MDD patients from Dataset B only. After the removal of one outlier, neither the association between spindle amplitude and the number of episodes nor with depression severity or outcome was significant (*p*s > .4), nor was the association between spindle density and depression severity, outcome, or number of episodes (*p*s > .2).

## Discussion

We aimed to systematically and comprehensively map non-REM sleep alterations of MDD patients against healthy controls. We explored the influence of non-REM sleep alterations on procedural memory consolidation and their relation to depression severity and outcome. Overall, no major alterations in non-REM sleep macrostructure were found in patients compared to controls. In contrast, a higher spindle amplitude was found in unmedicated patients compared to controls, whereas in medicated patients, longer SWs with lower amplitude and a more dispersed SW-spindle coupling were found. Overnight procedural memory consolidation was impaired only in medicated patients and was associated with lower sleep spindle density.

### Sleep macrostructure

Our reported changes in sleep architecture confirm earlier findings of REM sleep being reduced and delayed [4], as well as of lowered sleep efficiency, expressed in higher proportions of N1 and increased wake periods after sleep onset. Importantly, we only find these changes in MDD patients when medicated, not when unmedicated. Possibly, REM sleep alterations in the unmedicated MDD sample (Dataset B) were absent because these changes need time to manifest: these patients were relatively young and had the lowest number of depressive episodes beforehand. Alternatively, these changes are driven by the antidepressant medication, as they mirror well-described REM-suppressing effects expected by the antidepressants they received [15,42,43]. Unmedicated patients showed higher amounts of wake after sleep onset and worse subjective sleep quality compared to controls, which did not dissipate after short-term medication intake on the 7-day follow-up. However, we found hints of reduced arousal during sleep, expressed as reductions of alpha-band activity during non-REM [44, 45], only after prolonged medication in patients (i.e. after 28 days, Dataset C). This suggests that medication could objectively reduce night-time arousal, but does not subjectively increase sleep quality in the short term.

In contrast, changes in non-REM architecture were not as obvious in our samples. Only in pooled samples, a shorter SWS duration and its prolonged onset time became apparent in the medicated patients. Although a large proportion of the drugs our patients received are known to boost (e.g. certain types of TCA) or tend to increase SWS duration (e.g. SSRIs or Bupropion; [43], they seemed ineffective in doing so in our study. In our younger sample (Dataset B), lighter non-REM sleep (i.e. N2) increased under medication in the one-week follow-up. Surprisingly, although SWA was reduced in this follow-up, this did not manifest in reduced SWS time. This finding suggests that alterations in SWS duration may only be robust in larger samples.

### Slow-wave characteristics

A more detailed spectral analysis revealed that the reductions in SWS and SWA of medicated patients were specific to the upper delta band range (1–4 Hz). Recently, increases in this band range were revealed to be indicative for proper non-REM initialization and homeostatic, restorative processes, while lower bands (< 1 Hz), constituting the main activity used for scoring SWS, were not [46]. In line also with patients’ subjective sleep quality reports, this suggests that our medicated patients lacked such restorative and homeostatic features of non-REM sleep. In fact, general reductions in SWS and SWA have previously been reported in many MDD cohorts [5] even in younger samples that were unmedicated after a 2-week drug clearance [6, 47]. However, our unmedicated MDD patients did not show such reductions in delta/SWA bands or SWS, although they reported the lowest subjective sleep quality. Age-related SWS/SWA decline is unlikely to explain this [48], but depression history might. Our young unmedicated sample had markedly lower numbers of previous episodes than previous reports [6, 47] and we could link more depressive episodes to lower SW amplitudes in our samples. This also fits well with the null or opposite findings in samples with markedly lower depression severity [8, 9] or smaller sample size [7]. Again such SWA/delta band reductions in our medicated patients also mirror earlier reports of acute REM-suppressing effects of SSRI paroxetine [49]. However, no such reductions have been reported with other SSRI types such as trazodone, citalopram, fluvoxamine, or paroxetine [50–53]. Beta band activity, related to antidepressant drug use [49] or arousal in general [44], was altered in medicated and unmedicated state as well but, we could not attribute them consistently to any patient state. Taken together, it might thus be that typical MDD medication induces detrimental changes to REM and non-REM sleep that are long-lasting, i.e. their effects on sleep lasted longer than drug clearance periods of patients with a prior history of depression and medication (cf. [6]).

### Spindle characteristics

Interestingly, non-REM spindle band activity was increased independent of medication intake in the young patient group only (Dataset B). This was also reflected in intensified individual spindles (i.e. higher amplitude) of these patients. This pattern marked the most specific difference with controls. While reductions in spindle amplitude have previously been associated with sleep deprivation [54], this seems to be unlikely the case in our sample, given that our unmedicated sample spent more time awake after sleep onset, which counters to what is expected after sleep deprivation the previous night. In addition, one can speculate on the influence of a disturbed circadian rhythm in MDD patients compared to healthy [55], which are known to regulate spindle amplitude in younger (< 40 years) adults [56]. Unfortunately, our datasets did not include any measurements on circadian rhythm, therefore this remains to be directly tested. In contrast, our patients with the longest time on medication (i.e. Dataset A and Dataset C 28-day medication follow-up) showed lowered spindle density. This reduction was absent after combining Dataset A with the short-term medication (i.e. 7-day follow-up) of Dataset B and C against respective controls. Conversely, younger patients with no medication or after only 1-week of medication failed to show reductions in spindle density. Together this pattern of results seems to suggest that long-term, but not short-term medication lowers spindle density. This is however surprising, given that most patients were prescribed with antidepressants known to increase sleep spindle density after first use (e.g. SNRI, and some TCA) or leaving them unaffected (SSRI; [42]). It should however be noted that spindle density decreases with older age [57, 58]. Thus, the absence of spindle reductions in the markedly younger patients (in Dataset B) may suggest that age plays a synergistic and augmenting role in reducing spindles in depressed patients. In addition, again longer history of depression and concomitant long-term medication-induced changes might also account for such spindle density declines, as both were correlated as such. This can however only be answered by future studies including sufficiently drug-cleared or first-episode unmedicated and older patients.

### SW-spindle coupling

SW-spindles are thought to express the hippocampal-neocortical dialog necessary for memory consolidation processes during sleep [16]. Especially the accuracy and timing of SW-spindle coupling are vital for these processes [26,59,60]. We found that the number of spindles that lock to slow waves (SW-spindles), as well as their delay to the preceding slow-wave downstate, were unaltered in patient-control comparisons. However, the accuracy of this coupling (i.e. SW-spindle delay dispersion) worsened in all samples under medication. This was our strongest effect on sleep microstructure in the patient groups. Crucially, this mistiming of SW- spindles was not related to overnight memory procedural consolidation. Indeed, previous studies manipulating SW-spindles either through medication or transcranial stimulation, have shown only benefits for verbal and declarative memory consolidation [61, 62] but not procedural memory consolidation [63]. Thus, such changes might relate more to structural changes in hippocampal-cortical networks [26,59,64–66] related to degradation in patients than as a direct proxy for memory consolidation. However, we could not confirm any structural changes in hippocampal volumes in our MDD patients [21].

### Medication and psychiatric diagnosis

Different medication types show different effects on sleep features. Though not exhaustive, we subtyped patients by medication type across. We could recreate some, but not all, of the principal sleep alterations (e.g. reduced spindle density and SW amplitudes) and in part exclude mediating effects of demographic factors like age. If medication was driving all observed effects in our patients, then the supplementary explorative investigation could not capture this.

Other psychiatric populations were previously investigated on non-REM alterations, including for sleep spindle alterations. For example, several studies have specifically found robust spindle impairments in schizophrenia [67], related to cognitive performance, sleep-dependent procedural memory consolidation, and positive symptoms. Also, spindle amplitude has been found to be correlated with symptomology in schizophrenia and even suggested as a potential biomarker [68]. In addition, a preliminary but clear mechanistic understanding for this is in place, whereby a dysfunctioning thalamic reticular nucleus (TRN; where sleep spindles originate) is related to these spindle abnormalities, impaired sleep-dependent memory consolidation, as well as with symptomology in schizophrenia. Moreover, a recent study looked into the role of sleep spindles in bipolar patients in euthymic or stable mood [69]. Here, the authors found a reduction in spindle density in euthymic patients and link these results with the schizophrenia literature, such as overlap in terms of heritability between the disorders and responses to similar types of antipsychotics. Furthermore, the authors connected sensory gating deficits as seen in both bipolar as well as in schizophrenia patients with a dysfunctioning TRN. The contrast between the many investigations of sleep spindles in schizophrenia but few in MDD is surprising, especially since MDD is the more prevalent disease (World Health Organization, 2017). Here, we report similar sleep spindle density reductions in medicated MDD patients as also found in medicated schizophrenia patients, showing largely overlapping medication types including benzodiazepines, various antidepressants, and mood stabilizers.

Depression severity predicted reduced sleep spindles in medicated patients. Since this was not observed in the unmedicated sample, we speculate that decisions of physicians to prescribe a spindle-increasing vs decreasing drug might have indirectly related them to depression severity. In addition, diagnosis of depression is subjective and variable: it relies on self-reported symptoms of at least five out of nine symptoms (DSM-V) of which 256 combinations can be diagnosed as depression. Patients also differ in treatment response [70] and have been subdivided into so-called ‘biotypes’ by fMRI resting-state connectivity biomarkers in limbic and frontostriatal networks [71]. It is thus likely that subtypes of depression exist, but until now, have not been systematically distinguished. Therefore, given the variable definition of MDD and the observed variance within our cohorts and between other cohorts in the literature, one might speculate that the non-REM sleep alterations we reported may be found in certain biotypes of depression but not all. It thus remains open which impairments across psychiatric diseases can be explained by medication or single symptoms since many sleep studies have exclusively investigated in medicated patients and did not control well for potential patient subtypes. Future large-scale collaborative big-data sleep studies could be able to directly test this hypothesis using modern clustering techniques.

### Procedural memory consolidation

We also failed to confirm that procedural memory consolidation is impaired in our Dataset B as opposed to such previous reports in Dataset A [27]. Though procedures were comparable, the times of practice were not. Patients from Dataset B performed the task both in unmedicated and medicated state, providing a source of practice. Additionally, long-term medication might have induced partial psychomotor performance deficits [72, 73] affecting the procedural baseline levels, and thus also limit its subsequent consolidation. This was not observed in the group with short-term medication, which also showed higher baseline levels and may have benefited from previous training before medication onset. Further, age-depression interactions could also explain this discrepancy since group differences in overnight consolidation was previously observed in older individuals only [18]. Generally, overnight consolidation of procedural memory, similar to sleep spindle density, seems to decline with older age [74, 75] and may become even more apparent with additional psychiatric illness. In turn, older age inherently also leads to a higher chance of longer depressive episodes as well as more episodes in total. Possibly, lasting changes in procedural memory may occur in a cumulative manner, where under an acute state of depression, no observable changes have occurred yet. However, in our samples, the influence of the number of episodes on overnight consolidation performance was not significant. To tackle this question directly, future studies should look into long-term effects in bigger samples over time. In addition, when pooling available patients in the medicated state (Dataset A and B) they showed a stronger association between spindle density and overnight consolidation than controls. Thus, the actual lack of an association between spindle density and procedural memory consolidation in our healthy controls conflicts with previous reports (e.g.[76, 77]) although in line with similar attempts in MDD patients [78]. Furthermore, our findings could also be explained by the fact that our MDD sample showed more variance in spindle density overall, especially after combining the data of lower-performing patients of Dataset A with the higher-performing patients of Dataset B. In addition, this could point to underlying subtyping of MDD patients, where one subgroup represents high-functioning patients with intact sleep spindle densities and overnight consolidation ability whereas the other subgroup represents lower-functioning patients with impaired spindle density and consolidation. Lastly, in conflict with prior reports [20], procedural memory consolidation was also not predicted by hippocampal volumes (see supplementary materials for the analysis).

### Conclusions

Overall, our current explorative study suggests no clear spindle or non-REM or even REM deficits in depressed patients unless medicated. Only in the medicated sample did we find a consistent hampering of slow-wave, spindle activity, and reduced accuracy of SW-spindle coupling that aligned with procedural memory consolidation deficits in older but not younger medicated patients. In addition, procedural memory consolidation failed to consistently associate with SW-spindle mistiming and other non-REM features. Nevertheless, medication state and ensuing long-term effects in depressive patients seemed to be majorly but not exclusively associated with sleep alterations, and likely explains why sleep alterations overlap with other psychiatric populations in a medicated state. If sleep alterations map more closely to specific phenotypes of depression remains open. However, the medication-associated alterations are indicative of impaired memory and restorative processes during sleep but they alleviate little of the lost subjective sleep quality that patients report. This should caution physicians to weight the subtle but vital detriments of sleep features more carefully against more obvious benefits.

## Acknowledgments

Author Contributions. Leonore Bovy: Conceptualization, Methodology, Validation, Formal analysis, Data Curation, Writing - Original Draft, Visualization. Frederik D. Weber: Conceptualization, Methodology, Software, Validation, Formal analysis, Resources, Data Curation, Writing - Original Draft, Visualization. Indira Tendolkar: Supervision, Writing - Review & Editing. Guillen Fernandez: Supervision, Writing - Review & Editing. Michael Czisch: Investigation, Writing - Review & Editing. Axel Steiger: Writing - Review & Editing, Funding acquisition. Marcel Zeising: Investigation, Data Curation, Writing - Review & Editing. Martin Dresler: Conceptualization, Investigation, Data Curation, Writing - Original Draft, Supervision, Project administration, Funding acquisition.We thank Rathiga Varatheeswaran, Mariken C.R. Hoegen, and Marios Diamantopoulos for help in visual data pre-inspection of Dataset B.

