## supplementary materials for "Non-REM sleep in major depressive disorder"

**Results on combined data**

Here, we combined the three datasets by pooling all controls (*n* = 108) and all the patients in medicated state (*n* = 108). The patients after 7 days of medication were taken for Dataset C. All the main analysis were repeated on the combined data.

**Sleep architecture**. When combining the three datasets, the MDD patients spent a higher proportion in S1 sleep (ΔM = 4.33%, 95% CI [2.61%, 6.06%], t(198.29) = -4.96, p <.001) spent less time in SWS, ΔM = -2.8, 95% CI [-5.11, -0.49], t(208.86) = 2.39, p = .018. As expected, less proportion of sleep was spent in REM sleep (ΔM = -4.79%, 95% CI [-6.55%, 3.02%], t(189.75) = 5.35, p < .001) and patients took longer time to reach it compared to controls (ΔM = 97.58 min, 95% CI [76.95 min, 118.2 min], t(130.02) = 9.36, p <.001). Lastly, patients had less SWS (ΔM = -2.8 %, 95% CI [-5.11. %, -0.49 %], t(208.86) = -2.39, p = .018) and a later onset of SWS (ΔM = 7.85 min, 95% CI [2.36 min, 13.34 min], t(167.21)= -2.82, p = .005). See Table S5 for all results.

**Sleep spindles.** In the combined datasets, no group differences in spindle density nor in duration nor in amplitude were found. See Table S5 for all results.

**Slow waves.** Compared to controls, the medicated MDD patients had SW with lower SW amplitude (ΔM= -17.53 µV, 95% CI [-26.66 µV, -8.4 µV], *t*(213.28) = -3.78, *p* <.001), longer duration, (ΔM=0.04 s, 95% CI [0.01 s, 0.06 s], *t*(210.25) = 3.23, *p* = .001) and lower frequency, (ΔM= - 0.02 Hz 95% CI [-0.01 Hz, -0.04 Hz], *t*(211.22) = -3.17, *p* =.002). See Table S5 for all results.

**SW-spindles.** In the combined data, there were no group differences on the SW-spindle counts or differences in the mean spindle delay to SW of the coupling. MDD patients showed a greater delay dispersion – or spread around the mean, than controls (ΔM = 0.02 SD, 95% CI [0.01 SD, 0.03 SD], t(211.33) = 3.98, p <.001). See Table S5 for all results.

**Procedural memory.** When Dataset A and B (medicated sample) were combined on their memory performance (80 controls, 78 patients), there was a small effect at baseline after removal of one outlier, where MDD patients tapped less sequences correct than controls (ΔM = -1.29, 95% CI [-2.53, 0.05], $t(145.48) = 2.06$, $p = .042$). No differences in training effect were found (p = 0.23). However, MDD patients showed worse overnight consolidation than healthy controls after removal of 2 outliers (ΔM = -0.17, 95% CI [-0.25, -0.08], $t(122) = 3.94$, $p < .001$).

**Sleep parameters related to overnight memory consolidation.** When combining datasets A and B, an interaction effect between group (medicated MDD x controls) and spindle density on overnight consolidation re-emerged (after removal of two outliers: $b = 0.26$, 95% CI [0.03, 0.49], $t(148) = 2.22$, p = .028) which suggests a stronger association between spindle density and consolidation for patients ($r = 0.31$) than for controls ($r = 0.04$). No such interactions were found on SW parameters. In contrast, the significant interactions between the SW-spindle parameters in Dataset A could not be replicated in the combined dataset.

**Hippocampal volume**

In Dataset B, high-resolution structural magnetic resonance (MR) scans were acquired using a T1-weighted fast RF-spoiled gradient (FSPGR) sequence with the following parameters: TR = 6.18 ms; TE = 2.26 ms; flip angle = 12°; FOV = 256 mm; voxel resolution = 1 mm isotropic. MDD patients were in unmedicated state at the time of the MR scan. Data was available for 17 controls and 31 unmedicated patients. Data was available for 31 unmedicated patients and 17 controls. Subcortical structure segmentation was performed using the fsl (FMRIB Software Library) FIRST function. Next, volumetric analysis of the hippocampus was performed using fslstats based on standard labeling of the structures.

Given the previous mentioned literature on hippocampal size reductions in MDD, we wanted to explore if we could replicate these findings in our sample and correlate hippocampal size with overnight consolidation performance. Neither left or right, nor the average hippocampal volume (in $mm^3$) differed between the unmedicated MDD patients and controls ($p > 0.5$). An interaction between group and average hippocampal size on spindle frequency was found ($b = 0.00$, $t(44) = 2.36$, $p = .023$), suggesting a stronger negative correlation between hippocampal volume and spindle frequency in controls ($r = -0.67$, $p = 0.003$) than in patients ($r = -0.32$, $p = 0.076$). In addition, an interaction between group and average hippocampal size on delay between SW and spindles was found, ($b = 0.00$, $t(44) = 2.48$, $p = .017$). Here, controls show a moderate correlation between delay and hippocampal size ($r = 0.67$, $p = 0.004$), whereas patients do less so ($r = 0.16$, $p = 0.4$). See figure S1. Other sleep parameters nor overnight consolidation performance were correlated with hippocampal volume in either of the groups.

### Figure Captions

**Figure S1:** Hippocampal volume in Dataset B. (**A**) Hippocampal volume was stronger negatively correlated with spindle frequency in controls than in MDD patients. (**B**) Hippocampal volume was stronger correlated with mean delay between spindles and SW in controls than in MDD patients.

**Figure S2:** Behavioral results of finger tapping test of Dataset A and Dataset B combined. ***(A)*** Amount of correctly tapped sequences of first 30-second run. Medicated patients taped less sequences correct than Controls. (**B**) Percentage change score between the first run and the mean of the last three runs. (**C**) Percentage change score between the mean of three test runs after sleep in the morning and the mean of the last three training runs before sleep. Medicated patients perform worse after sleep than Controls. Data depicted like in Figures in the main text. Significances for two-group comparisons in asterisks (*, $p<.05$; ***, $p<.001$).

**Figure 1**

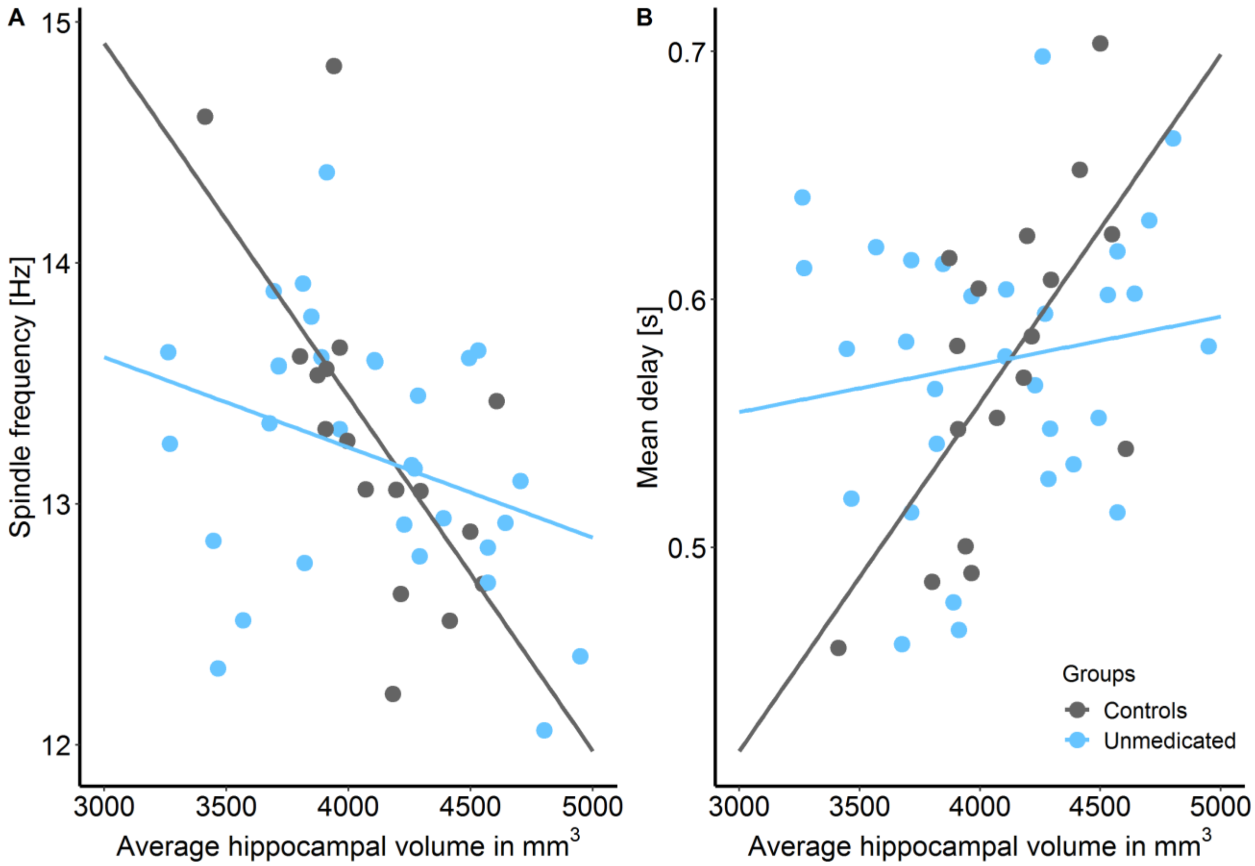
A
15
14
13
12
Spindle frequency [Hz]
3000
3500
4000
4500
5000
Average hippocampal volume in mm3
B
0.7
0.6
0.5
Mean delay [s]
3000
3500
4000
4500
5000
Average hippocampal volume in mm3
Groups
Controls
Unmedicated

**Figure S2**

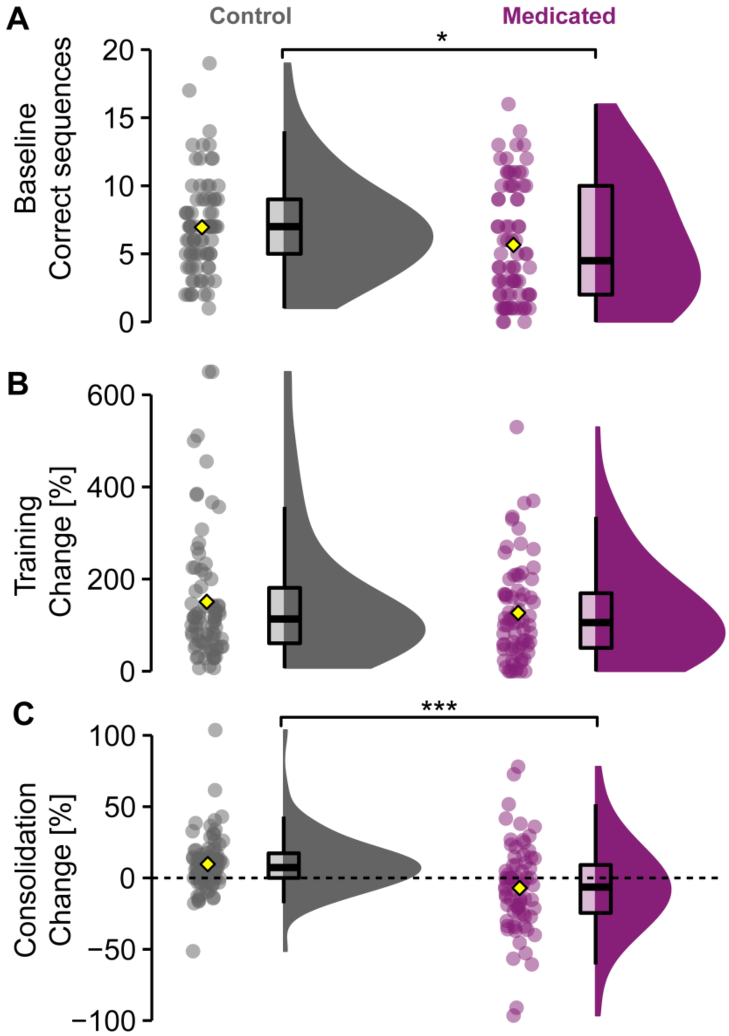
A
Control
Medicated
*
Baseline
Correct sequences
20
15
10
5
0
B
600
400
200
0
Training
Change [%]
C
***
100
50
0
−50
−100
Consolidation
Change [%]

**Table S1**

Sleep architecture statistical tests

| | Dataset A | Dataset B | | |
|---|---|---|---|---|
| | Medicated vs. controls | Unmedicated vs. controls | Unmedicated vs. medicated | Medicated vs. controls |
| N1[%] | **95% CI [2.50%, 8.32%], $t$(63.93) = 3.71, $p$ <.001, $BF_{10}$ = 68.92** | 95% CI [-2.7%, 3.02%], $t$(74.14) = 0.11, $p$ =.913, $BF_{10}$ = 0.24 | **95% CI [0.95%, 5.89%], $t$(37) = 2.8, $p$ =.008, $BF_{10}$ = 1.99** | **95% CI [0.71%, 6.43%], $t$(74.13) = 2.49, $p$ =.015, $BF_{10}$ = 3.3** |
| N2 [ %] | 95% CI [-0.53%, 8.74%], $t$(73.74) = 1.77, $p$ =.082, $BF_{10}$ = 0.89 | 95% CI [-5.2%, 1.39%], $t$(74.87) = -1.15, $p$ =.254, $BF_{10}$ = 0.42 | **95% CI [1.03%, 6.25%], $t$(37) = 2.83, $p$ =.008, $BF_{10}$ = 1.11** | 95% CI [-1.94%, 5.42%], $t$(69.7) = 0.94, $p$ =.35, $BF_{10}$ = 0.35 |
| SWS [%] | 95% CI [-7.23%, 1.19%], $t$(74.77) = -1.43, $p$ =.157, $BF_{10}$ = 0.56 | 95% CI [-4.99%, 2.02%], $t$(74.68) = 0.84, $p$ =.402, $BF_{10}$ = 0.32 | 95% CI [-3.28%, 0.32%], $t$(37) = 1.67, $p$ =.103, $BF_{10}$ = 0.32 | 95% CI [-6.4%, 0.47%], $t$(75.29) = 1.72, $p$ =.09, $BF_{10}$ = 0.84 |
| Non-REM [ %] | 95% CI [-3.4%, 5.57%], $t$(68.65) = 0.48, $p$ =.631, $BF_{10}$ = 0.26 | 95% CI [-7.21%, 0.43%], $t$(64.83) = -1.77, $p$ =.081, $BF_{10}$ = 0.93 | 95% CI [-0.88%, 5.2%], $t$(37) = 1.44, $p$ =.158, $BF_{10}$ = 0.35 | 95% CI [-5.16%, 2.7%], $t$(63.32) = -0.62, $p$ =.534, $BF_{10}$ = 0.28 |
| REM [ %] | **95% CI [-7.58%, -1.14%], $t$(63.70) = -2.71, p = .009, $BF_{10}$ = 5.21** | 95% CI [-2.36%, 2.82%], $t$(75.90) = -0.18, p = .862, $BF_{10}$ = 0.24 | **95% CI [-6.88%, -2.35%], $t$(37) = -4.39, p < .001, $BF_{10}$ = 31.99** | **95% CI [-7.65%, -2.23%], $t$(74.88) = -3.63, p = .001, $BF_{10}$ = 54.67** |
| WASO [ %] | 95% CI [-5.55%, 0.43%], $t$(72.24) = 1.71, $p$ =.092, $BF_{10}$ = 0.82 | **95% CI [0.94%, 6.07%], $t$(51.64) = 2.74, $p$ =.008, $BF_{10}$ = 6.26** | 95% CI [-3.16%, 1.48%], $t$(37) = -0.73, $p$ =.468, $BF_{10}$ = 0.27 | **95% CI [0.45%, 4.88%], $t$(57.12) = 2.41, $p$ =.019, $BF_{10}$ = 2.94** |
| TST [min] | 95% CI [-9.72min, 12.62min], $t$(75.71) = 0.26, $p$ =.797, $BF_{10}$ = 0.24 | 95% CI [-13.74min, 6.42min], $t$(52.24) = -0.73, $p$ =.47, $BF_{10}$ = 0.3 | 95% CI [-8.82min, 9.88min], $t$(37) = 0.11, $p$ =.91, $BF_{10}$ = 0.24 | 95% CI [-9.51min, 3.25min], $t$(74.14) = -0.98, $p$ =.331, $BF_{10}$ = 0.36 |
| Sleep onset [min] | 95% CI [-7.79min, 5.82min], $t$(74.05) = 0.29, $p$ =.774, $BF_{10}$ = 0.24 | 95% CI [-2.17min, 18.92min], $t$(44.08) = 1.6, $p$ =.117, $BF_{10}$ = 0.74 | 95% CI [-13.97min, 7.74min], $t$(37) = 0.58, $p$ =.564, $BF_{10}$ = 0.27 | 95% CI [-0.17min, 10.7min], $t$(66.5) = 1.93, $p$ =.058, $BF_{10}$ = 1.19 |
| SWS onset [min] | 95% CI [-1.25min, 21.5min], $t$(65.03) = 1.78, $p$ =.08, $BF_{10}$ = 0.91 | 95% CI [-0.49min, 17.17min], $t$(45.53) = 1.9, $p$ =.064, $BF_{10}$ = 1.18 | 95% CI [-9.95min, 9.12min], $t$(36) = 0.09, $p$ =.93, $BF_{10}$ = 0.24 | 95% CI [-0.35min, 15.48min], $t$(46.58) = 1.92, $p$ =.061, $BF_{10}$ = 1.26 |
| REM onset [min] | **95% CI [66.40min, 139.97min], $t$(45.48) = 5.65, $p$ <.001, $BF_{10}$ > 100** | 95% CI [-5.82min, 37.52min], $t$(62.64) = 1.46, $p$ =.149, $BF_{10}$ = 0.6 | **95% CI [36.56min, 101.67min], $t$(37) = 4.3, $p$ < .001, $BF_{10}$ > 100** | **95% CI [53.2min, 116.72min], $t$(48.16) = 5.38, $p$ < .001, $BF_{10}$ > 100** |

Non-REM is defined as the combination of N2 and SWS without N1. Statistical values represent a 95% percent confidence interval of the mean, a t-statisic and corresponding p-value (alpha = 0.05). In addition, Bayes factors are reported, where $BF_{10}$ <= 1 quantifies relative evidence in favor of the null hypothesis (H0), while a $BF_{10}$ > 1 quantifies relative evidence for the alternative hypothesis (H1). $BF_{10}$ values can be interpreted as either anecdotal (1-3), moderate (3-10), strong (10-30), very strong (30-100) or extreme (>100) evidence for H1.

**Table S1**.

Sleep architecture statistical tests – continued

| | Dataset C | | |
|---|---|---|---|
| | Medicated 7d vs. controls | Medicated 7d vs. Medicated 28d | Medicated 28d vs. controls |
| N1[%] | **95% CI [-1.38%, 6.87%], $t$(42.89) = 3.03, $p$ =.004, $BF_{10}$ = 9.14** | 95% CI [-2.48%, 3.87%], $t$(29) = 0.45, $p$ =.656, $BF_{10}$ = 0.28 | **95% CI [1.69%, 7.96%], $t$(37.45) = 3.11, $p$ =.003, $BF_{10}$ = 7.95** |
| N2 [ %] | 95% CI [-4.39%, 6.41%], $t$(52.08) = 0.38, $p$ =.709, $BF_{10}$ = 0.28 | 95% CI [-6.08%, 4.57%], $t$(29) = -0.29, $p$ =.775, $BF_{10}$ = 0.27 | 95% CI [-4.38%, 4.9%], $t$(55.95) = 0.11, $p$ =.911, $BF_{10}$ = 0.2 |
| SWS [%] | 95% CI [-6.48%, 2.37%], $t$(54.74) = 0.93, $p$ =.357, $BF_{10}$ = 0.38 | 95% CI [-4.4%, 5.48%], $t$(29) = 0.22, $p$ =.826, $BF_{10}$ = 0.27 | 95% CI [-6.1%, 3.07%], $t$(53.7) = 0.66, $p$ =.51, $BF_{10}$ = 0.24 |
| Non-REM [ %] | 95% CI [-5.25%, 3.17%], $t$(54.42) = -0.5, $p$ =.622, $BF_{10}$ = 0.29 | 95% CI [-5.03%, 4.61%], $t$(29) = -0.09, $p$ =.928, $BF_{10}$ = 0.26 | 95% CI [-5.25%, 3.17%], $t$(54.42) = -0.5, $p$ =.622, $BF_{10}$ = 0.23 |
| REM [ %] | **95% CI [-8.59%, 1.9%], $t$(46.82) = -3.16, p = .003, $BF_{10}$ = 12.43** | 95% CI [-2.91%, 4.89%], $t$(29) = -0.52, p = .608, $BF_{10}$ = 0.29 | **95% CI [-8.59%, -1.9%], $t$(46.82) = -3.16, p = .003, $BF_{10}$ = 2.66** |
| WASO [ %] | 95% CI [-1.48%, 5.13%], $t$(55.44) = 1.1, $p$ =.274, $BF_{10}$ = 0.44 | 95% CI [-5.04%, 1.41%], $t$(29) = -1.15, $p$ =.26, $BF_{10}$ = 0.49 | 95% CI [-1.48%, 5.13%], $t$(55.44) = 1.1, $p$ =.274, $BF_{10}$ = 0.2 |
| TST [min] | 95% CI [-12.3min, 6.25min], $t$(51.46) = -0.66, $p$ =.515, $BF_{10}$ = 0.32 | 95% CI [-5.34min, 9.54min], $t$(29) = 0.58, $p$ =.568, $BF_{10}$ = 0.3 | 95% CI [-12.3min, 6.25min], $t$(51.46) = -0.66, $p$ =.515, $BF_{10}$ = 0.2 |
| Sleep onset [min] | **95% CI [0.23min, 12.01min], $t$(44.64) = 2.09, $p$ =.042, $BF_{10}$ = 1.5** | 95% CI [-8.09min, 5.12min], $t$(29) = 0.46, $p$ =.65, $BF_{10}$ = 0.28 | **95% CI [0.23min, 12.01min], $t$(44.64) = 2.09, $p$ =.042, $BF_{10}$ = 0.57** |
| SWS onset [min] | 95% CI [-3.45min, 13.62min], $t$(49.73) = 1.2, $p$ =.237, $BF_{10}$ = 0.46 | 95% CI [-12.69min, 3.86min], $t$(28) = 1.09, $p$ =.28, $BF_{10}$ = 0.42 | 95% CI [-3.45min, 13.62min], $t$(49.73) = 1.2, $p$ =.237, $BF_{10}$ = 0.21 |
| REM onset [min] | **95% CI [64.5min, 149.02min], $t$(34.03) = 5.13, $p$ <.001, $BF_{10}$ > 100** | 95% CI [-64.44min, 51.03min], $t$(28) = 0.23, $p$ = .812, $BF_{10}$ = 0.27 | **95% CI [64.5min, 149.02min], $t$(34.03) = 5.13, $p$ < .001, $BF_{10}$ > 100** |

Non-REM is defined as the combination of N2 and SWS without N1. Statistical values represent a 95% percent confidence interval of the mean, a t-statisic and corresponding p-value (alpha = 0.05). In addition, Bayes factors are reported, where $BF_{10}$ <= 1 quantifies relative evidence in favor of the null hypothesis (H0), while a $BF_{10}$ > 1 quantifies relative evidence for the alternative hypothesis (H1). $BF_{10}$ values can be interpreted as either anecdotal (1-3), moderate (3-10), strong (10-30), very strong (30-100) or extreme (>100) evidence for H1.

**Table S2**

Overview of all sleep parameters of all three datasets (mean ± SE)

| | Dataset A | | Dataset B | | | Dataset C | | |
|---|---|---|---|---|---|---|---|---|
| | Controls | Medicated | Controls | Unmedicated | Medicated 7d | Controls | Medicated 7d | Medicated 28d |
| **Sleep spindles** | | | | | | | | |
| Density [/epoch] | 2.12 ± 0.06/**2.16 ± 0.05** | **1.94 ± 0.07**$^{*}$ | 2.34 ± 0.06 | 2.4 ± 0.06 | 2.42 ± 0.06 | **2.32 ± 0.07** | 2.15 ± 0.1 | **1.94 ± 0.11**$^{**}$ |
| Count | 1184 ± 43 | 1116 ± 63 | 1356 ± 46 | 1308 ± 58 | 1337 ± 56 | **1354 ± 55** | 1221 ± 70 | **1118 ± 77**$^{*}$ |
| Amplitude [µV] | 31.3 ± 1.41 | 28.8 ± 1.41 | **22.4 ± 0.85** | **25.2 ± 0.92**$^{\#}$ | **25.2 ± 0.92**$^{*}$ | 33.9 ± 1.6 | 31.6 ± 1.65 | 30.9 ± 1.62 |
| Frequency [Hz] | 13.1 ± 0.07 | 13.2 ± 0.1 | 13.3 ± 0.1 | 13.2 ± 0.09 | 13.1 ± 0.09 | 13.3 ± 0.1 | 13.1 ± 0.16 | 13.1 ± 0.15 |
| Duration [ms] | 771 ± 8 | 769 ± 9 | 784 ± 9 | 800 ± 8 | 808 ± 8 | **788 ± 10** | 771 ± 9 | **754 ± 10**$^{*}$ |
| **Slow waves** | | | | | | | | |
| Density [/epoch] | 1.42 ± 0.06 | 1.48 ± 0.07 | 1.51 ± 0.05 | 1.52 ± 0.06 | 1.49 ± 0.07 | 1.48 ± 0.07 | 1.4 ± 0.07 | 1.5 ± 0.08 |
| Count | 795 ± 40 | 846 ± 50 | 879 ± 38 | 841 ± 53 | 833 ± 54 | 870 ± 53 | 805 ± 51 | 879 ± 62 |
| Amplitude [µV] | **163 ± 5.25** | **136 ± 4.93**$^{***}$ | 148 ± 4.62 | 146 ± 5.43 | 145 ± 5.57 | **166 ± 6.74** | **143 ± 7.42**$^{*}$ | **146 ± 6.77**$^{*}$ |
| Frequency [Hz] | **0.84 ± 0.01** | **0.8 ± 0.01**$^{***}$ | 0.77 ± 0.01 | **0.77 ± 0.01** | **0.76 ± 0.01**$^{+}$ | 0.84 ± 0.01 | 0.81 ± 0.01 | 0.81 ± 0.01 |
| Duration [ms] | **1200 ± 8** | **1260 ± 12**$^{***}$ | 1300 ± 11 | **1310 ± 11** | **1330 ± 12**$^{+}$ | 1200 ± 10 | 1240 ± 17 | 1230 ± 15 |
| **SW-spindles** | | | | | | | | |
| Count | 90 ± 8 | 81 ± 6 | 106 ± 8 | 94 ± 9 | 92 ± 9 | 88 ± 8 | 80 ± 7 | 88 ± 9 |
| Delay [ms] | 540 ± 10 | 551 ± 10 | 556 ± 1 | 573 ± 9 | 576 ± 9 | 519 ± 20 | 492 ± 30 | 500 ± 23 |
| Delay dispersion [sd] | **0.215 ± 0.006** | **0.238 ± 0.007**$^{*}$ | **0.24 ± 0.006** | 0.253 ± 0.005 | **0.258 ± 0.007**$^{+}$ | **0.206 ± 0.006** | **0.24 ± 0.009**$^{**}$ | **0.235 ± 0.007**$^{*}$ |
| Spindle amplitude [µV] | 32.2 ± 1.17 | 29.5 ± 1.52 | **22.7 ± 0.88** | 25 ± 1.03 | **25.7 ± 0.94**$^{*}$ | 33.2 ± 1.51 | 31.9 ± 1.62 | 30.8 ± 1.68 |
| Spindle frequency [Hz] | 13 ± 0.07 | 13.1 ± 0.1 | 13.3 ± 0.1 | 13.1 ± 0.09 | 13.1 ± 0.1 | 13.2 ± 0.12 | 13 ± 0.15 | 13 ± 0.15 |
| Spindle duration [ms] | 706 ± 10 | 703 ± 10 | 711 ± 10 | 719 ± 10 | 710 ± 10 | **680 ± 10** | **710 ± 10**$^{*}$ | **690 ± 10**$^{\#}$ |
| SW amplitude [µV] | **164 ± 6.52** | **137 ± 5.66**$^{**}$ | 150 ± 5.53 | 153 ± 6.44 | 150 ± 6.64 | **171 ± 7.05** | **143 ± 7.65**$^{*}$ | **148 ± 6.82**$^{*}$ |
| SW duration [ms] | **1210 ± 10** | **1280 ± 20**$^{**}$ | 1320 ± 10 | 1340 ± 10 | 1340 ± 10 | **1190 ± 10** | **1250 ± 20**$^{**}$ | **1240 ± 20**$^{*}$ |
| Δ spindle amplitude [µV] | 0.551 ± 0.292 | 0.213 ± 0.386 | 0.249 ± 0.217 | -0.182 ± 0.192 | -0.054 ± 0.19 | -0.343 ± 0.353 | 0.312 ± 0.274 | 0.399 ± 0.207 |
| Δ spindle frequency [Hz] | -0.068 ± 0.013 | -0.07 ± 0.016 | **-0.017 ± 0.013** | **-0.03 ± 0.014** | **-0.068 ± 0.017**$^{*/+}$ | -0.054 ± 0.018 | -0.07 ± 0.17 | -0.024 ± 0.017 |
| Δ spindle duration [ms] | -0.072 ± 0.01 | -0.073 ± 0.011 | **-0.083 ± 0.009** | -0.096 ± 0.011 | **-0.109 ± 0.009***| **-0.113 ± 0.009** | **-0.075 ± 0.011**$^{**}$ | **-0.079 ± 0.011**$^{*}$ |
| Δ SW amplitude [µV] | 6.29 ± 1.94 | 6.35 ± 1.82 | 7.3 ± 1.39 | 7.06 ± 1.38 | 9.04 ± 1.27 | 9.94 ± 1.82 | 5.81 ± 1.37 | 7.68 ± 1.5 |
| Δ SW duration [ms] | 0.003 ± 0.009 | 0.018 ± 0.012 | 0.028 ± 0.008 | 0.018 ± 0.008 | 0.034 ± 0.01 | **-0.011 ± 0.009** | **0.016 ± 0.009*** | 0.003 ± 0.008 |

*Note* Non-REM is defined as the combination of N2 and SWS without N1. Epoch represents 30 seconds. Different symbols are used for indicating statistical comparisons within the datasets that are significant (highlighted in bold): differences between controls and medicated patients in Dataset A, B and C use asterisks (*, p <0.05; **, p<0.01; ***, p<0.001), between controls and unmedicated patients in Dataset B use hashes (#, p <0.05; ##, p<0.01; ###, p<0.001), and within patients for their follow-ups (i.e. Dataset B unmedicated vs. medicated 7d or Dataset C 7d medicated vs. 28d medicated patients) use pluses (+, p <0.05; ++, p<0.01; +++, p<0.001). Δ refers to difference between coupled and uncoupled spindles or slow waves (SW).

**Table S3**

Overview of all sleep parameters statistics

| | Dataset A | | Dataset B | |
|---|---|---|---|---|
| | Medicated vs. controls | Unmedicated vs. controls | Unmedicated vs. medicated | Medicated vs. controls |
| **Sleep spindles** | | | | |
| Density [/epoch] | **95% CI [0.4, 0.05], $t(70.79) = -2.58$, $p = .012$, $BF_{10} = 1.23$** | 95% CI [-0.11, 0.23], $t(75.73) = 0.73$, $p = .467$, $BF_{10} = 0.3$ | 95% CI [-0.04, 0.07], $t(37) = 0.64$, $p = .529$, $BF_{10} = 0.24$ | 95% CI [-0.08, 0.24], $t(74.58) = 0.98$, $p = .331$, $BF_{10} = 0.35$ |
| Count | 95% CI [-219.94, 84.29], $t(69.43) = -0.89$, $p = .377$, $BF_{10} = 0.33$ | 95% CI [-196.13, 98.93], $t(71.44) = -0.66$, $p = .513$, $BF_{10} = 0.28$ | 95% CI [-54.71, 113.32], $t(37) = 0.71$, $p = .484$, $BF_{10} = 0.25$ | 95% CI [-163.68, 125.08], $t(72.53) = -0.27$, $p = .791$, $BF_{10} = 0.24$ |
| Amplitude [µV] | 95% CI [-6.46 µV, 1.48 µV], $t(78) = -1.25$, $p = .215$, $BF_{10} = 0.47$ | **95% CI [0.27 µV, 5.26 µV], $t(75.06) = 2.21$, $p = .03$, $BF_{10} = 1.9$** | 95% CI [-0.82 µV, 0.82 µV], $t(37) < 0.001$, $p = .998$, $BF_{10} = 0.24$ | **95% CI [0.28 µV, 5.25 µV], $t(75.12) = 2.22$, $p = .03$, $BF_{10} = 1.92$** |
| Frequency [Hz] | 95% CI [-0.19 Hz, 0.28 Hz], $t(70.85) = 0.4$, $p = .69$, $BF_{10} = 0.25$ | 95% CI [-0.39 Hz, 0.13 Hz], $t(74.35) = 0.99$, $p = .327$, $BF_{10} = 0.36$ | 95% CI [-0.14 Hz, 0.05 Hz], $t(37) = 1.04$, $p = .305$, $BF_{10} = 0.25$ | 95% CI [-0.45 Hz, 0.09 Hz], $t(75.51) = 1.31$, $p = .194$, $BF_{10} = 0.49$ |
| Duration [ms] | 95% CI [-0.03ms, 0.02ms], $t(77.2) = -0.23$, $p = .821$, $BF_{10} = 0.24$ | 95% CI [-0.01ms, 0.04s], $t(75.96) = 1.24$, $p = .218$, $BF_{10} = 0.46$ | 95% CI [0.00ms, 0.02s], $t(37) = 1.59$, $p = .12$, $BF_{10} = 0.3$ | 95% CI [-0.00ms, 0.05s], $t(75.88) = 1.98$, $p = .051$, $BF_{10} = 1.25$ |
| **Slow waves** | | | | |
| Density [/epoch] | 95% CI [-0.11, 0.23], $t(76.85) = 0.7$, $p = 0.486$, $BF_{10} = 0.29$ | 95% CI [-0.18, 0.16], $t(73.09) = 0.11$, $p = 0.916$, $BF_{10} = 0.24$ | 95% CI [-0.12, 0.05], $t(37) = 0.8$, $p = .431$, $BF_{10} = 0.25$ | 95% CI [-0.2, 0.15], $t(71.22) = 0.28$, $p = 0.782$, $BF_{10} = 0.24$ |
| Count | 95% CI [-75.71, 178.09], $t(74.5) = -0.8$, $p = .424$, $BF_{10} = 0.31$ | 95% CI [-165.97, 91.22], $t(67.72) = -0.58$, $p = .564$, $BF_{10} = 0.27$ | 95% CI [-81.79, 64], $t(37) = -0.25$, $p = .806$, $BF_{10} = 0.24$ | 95% CI [-178.1, 85.55], $t(66.32) = -0.7$, $p = .486$, $BF_{10} = 0.29$ |
| Amplitude [µV] | **95% CI [-41.38 µV, -12.69 µV], $t(77.69) = -3.75$, $p < .001$, $BF_{10} = 77.42$** | 95% CI [-16.34 µV, -12.06 µV], $t(67.72) = -0.3$, $p = .765$, $BF_{10} = 0.24$ | 95% CI [-4.62 µV, 1.13 µV], $t(37) = 1.23$, $p = .226$, $BF_{10} = 0.24$ | 95% CI [-18.31 µV, 10.54 µV], $t(72.71) = -0.54$, $p = .593$, $BF_{10} = 0.27$ |
| Frequency [Hz] | **95% CI [0.02 Hz, 0.06 Hz], $t(72.07) = -3.77$, $p < .001$, $BF_{10} = 80.42$** | 95% CI [-0.02 Hz, 0.01 Hz], $t(76) = -0.35$, $p = .731$, $BF_{10} = 0.25$ | **95% CI [-0.02 Hz, 0.00 Hz], $t(37) = -2.33$, $p = .025$, $BF_{10} = 0.39$** | 95% CI [-0.03 Hz, 0.01 Hz], $t(75.33) = -1.4$, $p = .167$, $BF_{10} = 0.55$ |
| Duration [ms] | **95% CI [0.03ms, 0.09ms], $t(68.6) = -3.86$, $p < .001$, $BF_{10} > 100$** | 95% CI [0.03ms, 0.04ms], $t(75.97) = 0.3$, $p = .765$, $BF_{10} = 0.24$ | **95% CI [0.00ms, 0.03ms], $t(37) = 2.32$, $p = .026$, $BF_{10} = 0.41$** | 95% CI [-0.01ms, 0.06ms], $t(75.48) = 1.39$, $p = .170$, $BF_{10} = 0.54$ |
| **SW-spindles** | | | | |
| Count | 95% CI [-28.53, 12.28], $t(74.85) = -0.79$, $p = .43$, $BF_{10} = 0.31$ | 95% CI [-36.15, 12.99], $t(73.9) = -0.94$, $p = .351$, $BF_{10} = 0.34$ | 95% CI [-12.59, 8.56], $t(37) = -0.39$, $p = .702$, $BF_{10} = 0.24$ | 95% CI [-38.02, 10.84], $t(74.12) = -1.11$, $p = .271$, $BF_{10} = 0.4$ |

| | | | | |
|---|---|---|---|---|
| Delay [ms] | 95% CI [-0.02ms, 0.04ms], *t*(77.66) = 0.85, *p* =.395, $BF_{10}$ = 0.32 | 95% CI [-0.01ms, 0.04ms], *t*(75.49) = 1.28, *p* =.204, $BF_{10}$ = 0.47 | 95% CI [-0.01ms, 0.02ms], *t*(37) = 0.43, *p* =.67, $BF_{10}$ = 0.25 | 95% CI [-0.01ms, 0.05ms], *t*(75.8) = 1.53, *p* =.13, $BF_{10}$ = 0.64 |
| Delay dispersion [sd] | **95% CI [-0.04sd, 0.00sd], *t*(77.89) = 2.46, *p* =.016, $BF_{10}$ = 3.1** | 95% CI [-0.00sd, 0.03sd], *t*(74.65) = 1.7, *p* =.093, $BF_{10}$ = 0.8 | 95% CI [-0.01sd, 0.02sd], *t*(37) = 0.99, *p* =.331, $BF_{10}$ = 0.28 | **95% CI [0.00sd, 0.04sd], *t*(73.76) = 2.02, *p* =.047, $BF_{10}$ = 1.36** |
| Coupled spindle amplitude [µV] | 95% CI [-6.45 µV, 1.19 µV], *t*(73.39) = -1.37, *p* =.175, $BF_{10}$ = 0.52 | 95% CI [-0.43 µV, 4.95 µV], *t*(73.6) = 1.67, *p* =.098, $BF_{10}$ = 0.79 | 95% CI [-0.09 µV, 1.42 µV], *t*(37) = 1.78, *p* =.0.83, $BF_{10}$ = 0.26 | **95% CI [0.36 µV, 5.49 µV], *t*(75.32) = 2.27, *p* =.026, $BF_{10}$ = 2.13** |
| Coupled spindle frequency [Hz] | 95% CI [-0.18 Hz, 0.3 Hz], *t*(69.96) = -0.49, *p* = .624, $BF_{10}$ = 0.24 | 95% CI [-0.42 Hz, 0.12 Hz], *t*(74.49) = -1.1, *p* = .273, $BF_{10}$ = 0.29 | 95% CI [-0.19 Hz, 0.04 Hz], *t*(37) = -1.32, *p* = .195, $BF_{10}$ = 0.31 | 95% CI [-0.51 Hz, 0.06 Hz], *t*(76) = -1.56, *p* = .123, $BF_{10}$ = 0.24 |
| Coupled spindle duration [ms] | 95% CI [-0.03ms, 0.03ms], *t*(77.36) = -0.17, *p* = .864, $BF_{10}$ = 0.26 | 95% CI [-0.01ms, 0.03ms], *t*(75.65) = 0.68, *p* = .499, $BF_{10}$ = 0.4 | 95% CI [-0.02ms, 0.01ms], *t*(37) = -1.09, *p* = .282, $BF_{10}$ = 0.27 | 95% CI [-0.02ms, 0.02ms], *t*(74.93) = 0.07, *p* = .942, $BF_{10}$ = 0.67 |
| Coupled SW amplitude [µV] | **95% CI [-45.02 µV, -10.63 µV], *t*(76.47) = -3.22, *p* =.002, $BF_{10}$ = 17.96** | 95% CI [-13.64 µV, -20.17 µV], *t*(73.67) = 0.38, *p* =.701, $BF_{10}$ = 0.25 | 95% CI [-9.03 µV, -2.99 µV], *t*(37) = -1.02, *p* =.315, $BF_{10}$ = 0.25 | 95% CI [-16.96 µV, -17.46 µV], *t*(73.96) = 0.03, *p* =.977, $BF_{10}$ = 0.24 |
| Coupled SW duration [ms] | **95% CI [0.03ms, 0.12ms], *t*(71.75) = 3.37, *p* = .001, $BF_{10}$ = 26.7** | 95% CI [-0.02ms, 0.05ms], *t*(71.95) = -0.98, *p* = .330, $BF_{10}$ = 0.36 | 95% CI [-0.02ms, 0.03ms], *t*(37) = 0.46, *p* = .646, $BF_{10}$ = 0.25 | 95% CI [-0.01ms, 0.06ms], *t*(73.96) = 1.36, *p* = .178, $BF_{10}$ = 0.52 |
| Δ coupled- uncoupled spindle amplitude [µV] | 95% CI [-1.3 µV, 0.63 µV], *t*(72.69) = -0.7, *p* =.487, $BF_{10}$ = 0.29 | 95% CI [-1.01 µV, 0.15 µV], *t*(72.25) = 1.49, *p* =.141, $BF_{10}$ = 0.6 | 95% CI [-0.2 µV, 0.46 µV], *t*(37) = 0.79, *p* =.436, $BF_{10}$ = 0.26 | 95% CI [-0.88 µV, 0.27 µV], *t*(75.12) = 1.05, *p* =.297, $BF_{10}$ = 0.377 |
| Δ coupled- uncoupled spindle frequency [Hz] | 95% CI [-0.04 Hz, 0.04 Hz], *t*(74.18) = -0.05, *p* = 0.958, $BF_{10}$ = 0.23 | 95% CI [-0.05 Hz, 0.02 Hz], *t*(74.79) = -0.69, *p* = .49, $BF_{10}$ = 0.29 | **95% CI [-0.07 Hz, -0.00 Hz], *t*(37) = 2.06, *p* = .046, $BF_{10}$ = 0.83** | **95% CI [-0.09 Hz, -0.01 Hz], *t*(68.99) = -2.39, *p* = .02, $BF_{10}$ = 2.75** |
| Δ coupled- uncoupled spindle duration [ms] | 95% CI [-0.03ms, 0.03ms], *t*(76.94) = -0.07, *p* = .941, $BF_{10}$ = 0.23 | 95% CI [-0.04ms, 0.01ms], *t*(73.22) = -0.93, *p* = .355, $BF_{10}$ = 0.34 | 95% CI [-0.03ms, 0.00ms], *t*(37) = -1.68, *p* = .102, $BF_{10}$ = 0.36 | **95% CI [-0.05ms, 0.00ms], *t*(75.95) = -2.1, *p* = .039, $BF_{10}$ = 1.54** |
| Δ coupled- uncoupled SW amplitude [µV] | 95% CI [-5.24 µV, -5.35 µV], *t*(77.69) = 0.02, *p* =.984, $BF_{10}$ = 0.23 | 95% CI [-4.16 µV, 3.66 µV], *t*(75.97) = 0.13, *p* =.899, $BF_{10}$ = 0.24 | 95% CI [-0.79 µV, 4.77 µV], *t*(37) = 1.45, *p* =.156, $BF_{10}$ = 0.38 | 95% CI [-2.02µV, 5.49 µV], *t*(75.7) = 0.92, *p* =.36, $BF_{10}$ = 0.34 |
| Δ coupled- uncoupled SW duration [ms] | 95% CI [-0.02ms, 0.05ms], *t*(72.97) = 0.97, *p* = .336, $BF_{10}$ = 0.35 | 95% CI [-0.03ms, 0.01ms], *t*(75.89) = 0.83, *p* = .409, $BF_{10}$ = 0.32 | 95% CI [-0.01ms, 0.04ms], *t*(37) = 1.55, *p* = .13, $BF_{10}$ = 0.48 | 95% CI [-0.02ms, 0.03ms], *t*(72.91) = 0.53, *p* = .599, $BF_{10}$ = 0.27 |

Non-REM is defined as the combination of N2 and SWS without N1. Statistical values represent a 95% percent confidence interval of the mean, a t-statisic and corresponding p-value (alpha = 0.05). In addition, Bayes factors are reported, where $BF_{10}$ <= 1 quantifies relative evidence in favor of the null hypothesis (H0), while a $BF_{10}$ > 1 quantifies relative evidence for the alternative hypothesis (H1). $BF_{10}$ values can be interpreted as either anecdotal (1-3), moderate (3-10), strong (10-30), very strong (30-100) or extreme (>100) evidence for H1.

**Table S3**

Overview of all sleep parameters statistics – continued

| | Dataset C | | |
|---|---|---|---|
| | Medicated 7d vs. controls | Medicated 7d vs. medicated 28d | Medicated 28d vs. controls |
| | | Sleep spindles | |
| Density [/epoch] | 95% CI [-0.42, 0.08], $t$(52.67) = -1.38, $p$ = .175, $BF_{10}$ = 0.57 | 95% CI [-0.5, 0.08], $t$(29) = -1.45, $p$ = .157, $BF_{10}$ = 0.58 | **95% CI [-0.65, -0.11], $t$(49.21) = -2.79, $p$ = .007, $BF_{10}$ = 4.4** |
| Count | 95% CI [-311.94, 45.49], $t$(53.91) = -1.49, $p$ =.141, $BF_{10}$ = 0.66 | 95% CI [-293.72, 87.95], $t$(29) = -1.1, $p$ =.297, $BF_{10}$ = 0.4 | **95% CI [-425.49, -46.73], $t$(51.94) = -2.5, $p$ =.016, $BF_{10}$ = 2.58** |
| Amplitude [µV] | 95% CI [-6.86 µV, 2.33 µV], $t$(56) = -0.99, $p$ =.328, $BF_{10}$ = 0.4 | 95% CI [-4.13 µV, 2.59 µV], $t$(29) = -0.47, $p$ =.642, $BF_{10}$ = 0.27 | 95% CI [-7.59 µV, 1.52 µV], $t$(55.98) = -1.33, $p$ =.187, $BF_{10}$ = 0.44 |
| Frequency [Hz] | 95% CI [-0.56 Hz, 0.19 Hz], $t$(49.42) = -1.01, $p$ =.316, $BF_{10}$ = 0.4 | 95% CI [-0.46 Hz, 0.44 Hz], $t$(29) = -0.04, $p$ =.965, $BF_{10}$ = 0.26 | 95% CI [-0.56 Hz, 0.16 Hz], $t$(50.52) = -1.1, $p$ =.278, $BF_{10}$ = 0.34 |
| Duration [ms] | 95% CI [-0.04ms, 0.01s], $t$(55.58) = -1.27, $p$ =.208, $BF_{10}$ = 0.52 | 95% CI [-0.04ms, 0.01s], $t$(29) = -1.29, $p$ =.207, $BF_{10}$ = 0.5 | **95% CI [-0.06ms, -0.01s], $t$(55.98) = -2.44, $p$ =.018, $BF_{10}$ = 2.42** |
| | | Slow waves | |
| Density [/epoch] | 95% CI [-0.28, 0.12], $t$(55.73) = 0.78, $p$ = 0.437, $BF_{10}$ = 0.34 | 95% CI [-0.12, 0.34], $t$(29) = 0.96, $p$ = 0.347, $BF_{10}$ = 0.41 | 95% CI [-0.18, 0.24], $t$(55.82) = 0.28, $p$ = 0.783, $BF_{10}$ = 0.21 |
| Count | 95% CI [-211.05, 81.05], $t$(55.71) = -0.89, $p$ =.376, $BF_{10}$ = 0.37 | 95% CI [-89.23, 237.96], $t$(29) = 0.93, $p$ =.36,$BF_{10}$ = 0.38 | 95% CI [-152.99, 171.72], $t$(55.13) = 0.12, $p$ =.908, $BF_{10}$ = 0.2 |
| Amplitude [µV] | **95% CI [-43.38 µV, -3.23 µV], $t$(55.79) = -2.33, $p$ = .024, $BF_{10}$ = 2.38** | 95% CI [15.66 µV, 22.75 µV], $t$(29) = 0.38, $p$ = .709, $BF_{10}$ = 0.28 | **95% CI [-38.89 µV, -0.63 µV], $t$(55.95) = -2.07, $p$ = .043, $BF_{10}$ = 1.26** |
| Frequency [Hz] | 95% CI [-0.05 Hz, 0.01 Hz], $t$(48.25) = -1.59, $p$ =.119, $BF_{10}$ = 0.73 | 95% CI [-0.03 Hz, 0.03 Hz], $t$(29) = 0.02, $p$ =.981, $BF_{10}$ = 0.26 | 95% CI [-0.04 Hz, 0.00 Hz], $t$(52.15) = -1.73, $p$ =.089, $BF_{10}$ = 0.73 |
| Duration [ms] | 95% CI [-0.01ms, 0.07ms], $t$(45.98) = 1.75, $p$ =.087, $BF_{10}$ = 0.92 | 95% CI [-0.05ms, 0.04ms], $t$(29) = -0.11, $p$ =.915, $BF_{10}$ = 0.15 | 95% CI [-0.00ms, 0.07ms], $t$(50.10) = 1.83, $p$ =.074, $BF_{10}$ = 0.83 |
| | | SW-spindles | |
| Count | 95% CI [-31.17, 17.38], $t$(55.91) = -0.57, $p$ =.572, $BF_{10}$ = 0.3 | 95% CI [-20.5, 36.9], $t$(37) = 0.58, $p$ =.564, $BF_{10}$ = 0.3 | 95% CI [-26.43, 29.04], $t$(52.6) = 0.09, $p$ =.925, $BF_{10}$ = 0.2 |
| Delay [ms] | 95% CI [-0.02ms, 0.05ms], $t$(51.03) = 0.9, $p$ =.374, $BF_{10}$ = 0.37 | 95% CI [-0.02ms, 0.04ms], $t$(29) = -0.64, $p$ =.527, $BF_{10}$ = 0.3 | 95% CI [-0.03ms, 0.04ms], $t$(53.26) = 0.3, $p$ =763, $BF_{10}$ = 0.21 |
| Delay dispersion [sd] | **95% CI [0.01sd, 0.05sd], $t$(49.61) = 3.03, $p$ =.004, $BF_{10}$ = 9.42** | 95% CI [-0.03sd, 0.01sd], $t$(29) = -1.23, $p$ =.228, $BF_{10}$ = 0.39 | **95% CI [0.00sd, 0.04sd], $t$(54.5) = 2.28, $p$ =.026, $BF_{10}$ = 1.76** |
| Coupled spindle amplitude [µV] | 95% CI [-5.78 µV, 3.09 µV], $t$(55.94) = 0.61, $p$ =.546, $BF_{10}$ = 0.31 | 95% CI [-4.56 µV, 2.34 µV], $t$(29) = 0.66, $p$ =.516, $BF_{10}$ = 0.29 | 95% CI [-6.98 µV, -2.07 µV], $t$(55.75) = -1.09, $p$ =.281, $BF_{10}$ = 0.34 |

| | | | |
|---|---|---|---|
| Coupled spindle frequency [Hz] | 95% CI [-0.58 Hz, 0.17 Hz], *t*(51.27) = -1.09, *p* = 0.279, $BF_{10}$ = 0.43 | 95% CI [-0.39 Hz, 0.48 Hz], *t*(29) = 0.21, *p* = .838, $BF_{10}$ = 0.27 | 95% CI [-0.53 Hz, 0.21 Hz], *t*(51.6) = -0.86, *p* = .391, $BF_{10}$ = 0.22 |
| Coupled spindle duration [ms] | **95% CI [0.01ms, 0.04ms], *t*(54.77) = 2.53, *p* = .014, $BF_{10}$ = 3.48** | 95% CI [-0.04ms, 0.00ms], *t*(29) = -2.03, *p* = .052, $BF_{10}$ = 1.49 | 95% CI [-0.01ms, 0.02ms], *t*(55.79) = 0.42, *p* = .678, $BF_{10}$ = 0.28 |
| Coupled SW amplitude [µV] | **95% CI [-48.21 µV, -6.52 µV], *t*(55.88) = 2.63, *p* =.011, $BF_{10}$ = 4.29** | 95% CI [-15.67 µV, 25.27 µV], *t*(29) = 0.48, *p* =.635, $BF_{10}$ = 0.29 | **95% CI [-42.21 µV, -2.91 µV], *t*(55.73) = 2.3, *p* =.025, $BF_{10}$= 1.88** |
| Coupled SW duration [ms] | **95% CI [0.02ms, 0.011ms], *t*(50.29) = 2.8, *p* = .007, $BF_{10}$ = 5.8** | 95% CI [-0.04ms, 0.02ms], *t*(29) = -0.54, *p* = .593, $BF_{10}$ = 0.28 | **95% CI [0.01ms, 0.1ms], *t*(52.54) = 2.56, *p* = .013, $BF_{10}$= 2.88** |
| Δ coupled- uncoupled spindle amplitude [µV] | 95% CI [-0.24 µV, 1.55 µV], *t*(51.81) = 1.46, *p* =.149, $BF_{10}$ = 0.66 | 95% CI [-0.5 µV, 0.68 µV], *t*(29) = 0.3, *p* =.764, $BF_{10}$ = 0.27 | 95% CI [-0.08 µV, 1.57 µV], *t*(43.92) = 1.81, *p* =.077, $BF_{10}$ = 0.88 |
| Δ coupled- uncoupled spindle frequency [Hz] | 95% CI [-0.07 Hz, 0.03 Hz], *t*(55.6) = 0.65, *p* = .52, $BF_{10}$ = 5.14 | 95% CI [0.00 Hz, 0.09 Hz], *t*(29) = 2.01, *p* = .054, $BF_{10}$ = 1.2 | 95% CI [-0.02 Hz, 0.08 Hz], *t*(55.02) = 1.2, *p* = .234, $BF_{10}$ = 0.39 |
| Δ coupled- uncoupled spindle duration [ms] | **95% CI [0.01ms, 0.07ms], *t*(55.31) = 2.72, *p* = .009, $BF_{10}$ = 0.32** | 95% CI [-0.03ms, 0.02ms], *t*(29) = -0.37, *p* = .712, $BF_{10}$ = 0.27 | **95% CI [0.01ms, 0.06ms], *t*(55.24) = 2.43, *p* = .018, $BF_{10}$= 2.32** |
| Δ coupled- uncoupled SW amplitude [µV] | 95% CI [-8.71 µV, 0.45 µV], *t*(50.99) = 1.81, *p* =.076, $BF_{10}$ = 1.05 | 95% CI [-2.31 µV, 6.04 µV], *t*(29) = 0.91, *p* =.369, $BF_{10}$ = 0.37 | 95% CI [-7.01 µV, 2.47 µV], *t*(53.25) = -0.96, *p* =.341, $BF_{10}$ = 0.31 |
| Δ coupled- uncoupled SW duration [ms] | **95% CI [0.00ms, 0.05ms], *t*(55.84) = 2.12, *p* = .039, $BF_{10}$ = 1.67** | 95% CI [-0.03ms, 0.01ms], *t*(29) = -1.37, *p* = .181, $BF_{10}$ = 0.43 | 95% CI [-0.01ms, 0.04ms], *t*(54.75) = 1.14, *p* = .26, $BF_{10}$= 0.36 |

Non-REM is defined as the combination of N2 and SWS without N1. Statistical values represent a 95% percent confidence interval of the mean, a t-statisic and corresponding p-value (alpha = 0.05). In addition, Bayes factors are reported, where $BF_{10}$ <= 1 quantifies relative evidence in favor of the null hypothesis (H0), while a $BF_{10}$ > 1 quantifies relative evidence for the alternative hypothesis (H1). $BF_{10}$ values can be interpreted as either anecdotal (1-3), moderate (3-10), strong (10-30), very strong (30-100) or extreme (>100) evidence for H1.

**Table S4**

Sleep cycle durations split by non-REM and REM

| Dataset | Group | Cycle | N | Duration Cycle [min] median | Duration Cycle [min] mean | Duration Cycle [min] SE | Duration Non-REM [min] median | Duration Non-REM [min] mean | Duration Non-REM [min] SE | Duration REM [min] median | Duration REM [min] mean | Duration REM [min] SE |
|---|---|---|---|---|---|---|---|---|---|---|---|---|
| A | Controls | 1 | 40 | 87 | **91.72** | 4.89 | 70.75 | **75.44** | 4.85 | 15.75 | 16.29 | 1.37 |
| | | 2 | 40 | 99 | **100.76** | 3.14 | 78.75 | **80.78** | 2.64 | 19.75 | **19.99** | 1.59 |
| | | 3 | 39 | 114 | 114.51 | 4.71 | 84.5 | 83.38 | 3.4 | 30.5 | 31.13 | 2.9 |
| | | 4 | 33 | 99 | 102.64 | 5.15 | 69 | **71.17** | 4.26 | 28.5 | 31.47 | 2.58 |
| | | 5 | 15 | 72.5 | 77.73 | 4.8 | 58.5 | 56.1 | 2.66 | 14 | 21.63 | 4.01 |
| | | 6 | 2 | 65.75 | 65.75 | 18.75 | 54.75 | 54.75 | 13.75 | 11 | 11 | 5 |
| | Medicated | 1 | 40 | 169.75 | **196.24/*** ** | 17.31 | 149.75 | **177.04/*** ** | 17.58 | 16.75 | 19.2 | 2.21 |
| | | 2 | 37 | 132.5 | **133.73/*** ** | 7.01 | 92 | **101.08/** ** | 5.8 | 26 | **32.65/*** ** | 2.92 |
| | | 3 | 23 | 111 | 114.8 | 7.63 | 81 | 83.65 | 5.9 | 29.5 | 31.15 | 4.34 |
| | | 4 | 11 | 95 | 96.95 | 6.4 | 56.5 | **57.23/* ** | 3.6 | 30.5 | 39.73 | 9.03 |
| | | 5 | 4 | 82.25 | 78.38 | 19.16 | 54.75 | 49.62 | 9.47 | 21 | 28.75 | 13.14 |
| | | 6 | 3 | 85 | 78.17 | 10.53 | 55.5 | 55.17 | 7.51 | 24 | 23 | 11.85 |
| B | Controls | 1 | 40 | 96.25 | **103.36** | 6.65 | 84.25 | **89.44** | 5.76 | 12.75 | 13.93 | 1.9 |
| | | 2 | 40 | 101.75 | **108.24** | 3.95 | 82.25 | 86.38 | 3.44 | 21.75 | 21.86 | 2.05 |
| | | 3 | 40 | 104.75 | 107.03 | 3.65 | 80.25 | 80.6 | 2.85 | 25.25 | 26.43 | 2.43 |
| | | 4 | 37 | 87.5 | 96.89 | 3.56 | 66.5 | 67.58 | 2.75 | 30 | 29.31 | 2.41 |
| | | 5 | 17 | 64.5 | 71.53 | 5.84 | 47 | 48.94 | 4.18 | 17.5 | 22.59 | 3.49 |
| | | 6 | 4 | 38 | 50.38 | 15.6 | 24.25 | 39.25 | 17.2 | 8.5 | 11.12 | 4.62 |
| | | 7 | 2 | 27.75 | 27.75 | 8.25 | 19.75 | 19.75 | 0.75 | 8 | 8 | 7.5 |
| | Unmedicated | 1 | 38 | 102 | **122.86** | 10.3 | 87 | **105.28** | 9.19 | 17.5 | 17.58 | 1.72 |
| | | 2 | 37 | 119 | 119.89 | 6.5 | 85.5 | 93.19 | 5.53 | 24.5 | 26.7 | 2.55 |
| | | 3 | 33 | 102.5 | 105.03 | 4.72 | 76.5 | 79.95 | 4.72 | 24.5 | 25.08 | 1.55 |
| | | 4 | 28 | 85.5 | 85.54 | 5.34 | 62.25 | 59.71 | 3.09 | 19.75 | 25.82 | 3.47 |
| | | 5 | 14 | 81.5 | 73.54 | 8.05 | 52.5 | 46.46 | 4.78 | 26 | 27.07 | 4.38 |
| | | 6 | 5 | 48.5 | 51.4 | 12.04 | 44 | 47.2 | 10.96 | 3.5 | 4.2 | 1.57 |
| | | 7 | 1 | 25 | 25 | | 18.5 | 18.5 | | 6.5 | 6.5 | |
| | Medicated 7d | 1 | 38 | 185 | **192.7/***/ ###** | 16.12 | 169.5 | **172.53/*** /###** | 15.17 | 16.75 | 20.17 | 2.73 |
| | | 2 | 36 | 121.5 | **124.85/* ** | 7.23 | 99 | 101.08 | 6.48 | 22 | 23.76 | 2.4 |

| | | | | | | | | | | | | |
|---|---|---|---|---|---|---|---|---|---|---|---|---|
| | | 3 | 27 | 112 | 112.87 | 6.74 | 89 | 89.83 | 4.88 | 18.5 | 23.04 | 3.19 |
| | | 4 | 12 | 95.25 | 99.92 | 9.19 | 72.75 | 69.62 | 4.98 | 29 | 30.29 | 5.39 |
| | | 5 | *4* | 72.5 | 66.62 | 10.37 | 62.5 | 56.12 | 9.97 | 10.25 | *10.5*/** | 1.34 |
| | | 6 | *2* | 55.75 | 55.75 | 2.25 | 40 | 40 | 8.5 | 15.75 | 15.75 | 10.75 |
| | | 7 | *1* | 48.5 | 48.5 | | 44.5 | 44.5 | | 4 | 4 | |
| C | Controls | 1 | 28 | 96.75 | **97.11** | 7.29 | 76 | **79.16** | 5.91 | 15.25 | **17.95** | 2.11 |
| | | 2 | 28 | 98.25 | **105.45** | 5.16 | 74.25 | **82.64** | 4.18 | 20.75 | 22.8 | 2.72 |
| | | 3 | 28 | 104.5 | 104.07 | 3.51 | 74.75 | 77.96 | 3.09 | 27 | 26.11 | 1.85 |
| | | 4 | 26 | 105.5 | 102.65 | 3.07 | 71.5 | 72.75 | 2.42 | 29.75 | 29.9 | 2.3 |
| | | 5 | 13 | 85 | 85.62 | 5.26 | 69 | 65.81 | 4.66 | 16.5 | 19.81 | 3.54 |
| | Medicated 7d | 1 | 30 | 184.25 | **206.73/*** ** | 20.74 | 164.75 | **186.8/*** ** | 19.94 | 16.75 | 19.93 | 2.7 |
| | | 2 | 27 | 125.5 | 124.83 | 8.24 | 97 | 94 | 6.94 | 29.5 | 30.83 | 3.46 |
| | | 3 | 16 | 119.5 | 123.94 | 9.63 | 82.5 | 90.94 | 8.79 | 31.5 | 33 | 4.3 |
| | | 4 | 9 | 87 | **88.33/*** | 5.46 | 59 | 65.11 | 5.37 | 19 | 23.22 | 7.03 |
| | | 5 | 2 | 64.25 | 64.25 | 45.75 | 30.75 | 30.75 | 13.25 | 33.5 | 33.5 | 32.5 |
| | | 6 | 1 | 116.5 | 116.5 | | 52 | 52 | | 64.5 | 64.5 | |
| | Medicated 28d | 1 | 29 | 202 | **203.79/*** ** | 18.35 | 183 | **174.83/*** ** | 18.52 | 22.5 | **28.97/** ** | 3.48 |
| | | 2 | 27 | 121.5 | **132.44/*** | 8.91 | 96 | **105.11/*** | 7.3 | 26.5 | 27.33 | 3.79 |
| | | 3 | 21 | 101.5 | 97.4 | 9.26 | 73.5 | 71.38 | 8.09 | 17.5 | 26.02 | 4.21 |
| | | 4 | 10 | 90.5 | 89.7 | 11.54 | 60 | 62.15 | 9.71 | 21.5 | 27.55 | 7.96 |
| | | 5 | *4* | 58.25 | 61.25 | 15.03 | 29 | *32.25*/*** | 4.21 | 29.25 | 29 | 11.46 |
| | | 6 | *3* | 35 | 62 | 35.84 | 28.5 | 36 | 13.38 | 6.5 | 26 | 22.57 |
| **REM suppressing medication:** | | | | | | | | | | | | |
| A | Controls | 1 | 40 | 87 | **91.72** | 4.89 | 70.75 | **75.44** | 4.85 | 15.75 | 16.29 | 1.37 |
| | | 2 | 40 | 99 | **100.76** | 3.14 | 78.75 | **80.78** | 2.64 | 19.75 | **19.99** | 1.59 |
| | | 3 | 39 | 114 | 114.51 | 4.71 | 84.5 | 83.38 | 3.4 | 30.5 | 31.13 | 2.9 |
| | | 4 | 33 | 99 | 102.64 | 5.15 | 69 | **71.17** | 4.26 | 28.5 | 31.47 | 2.58 |
| | | 5 | 15 | 72.5 | 77.73 | 4.8 | 58.5 | 56.1 | 2.66 | 14 | 21.63 | 4.01 |
| | | 6 | 2 | 65.75 | 65.75 | 18.75 | 54.75 | 54.75 | 13.75 | 11 | 11 | 5 |
| | Medicated | 1 | 34 | 186 | **204.62/*** ** | 18.43 | 155.75 | **187.24/*** ** | 18.59 | 14.75 | 17.38 | 2.27 |
| | | 2 | 31 | 132.5 | **133.74/*** ** | 7.94 | 92 | **101.16/** ** | 6.66 | 26 | **32.58/** ** | 3.22 |
| | | 3 | 18 | 109 | 116.67 | 9.47 | 82.5 | 85.08 | 7.33 | 31 | 31.58 | 5.15 |
| | | 4 | 7 | 95 | 98.21 | 9.94 | 56.5 | **54.43/** ** | 3.95 | 30.5 | 43.79 | 12.85 |
| | | 5 | 3 | 65.5 | 65.17 | 19.63 | 56 | 48.33 | 13.27 | 9.5 | 16.83 | 7.84 |
| | | 6 | 3 | 85 | 78.17 | 10.53 | 55.5 | 55.17 | 7.51 | 24 | 23 | 11.85 |

| | | | | | | | | | | | | |
|---|---|---|---|---|---|---|---|---|---|---|---|---|
| B | Controls | 1 | 40 | 96.25 | **103.36** | 6.65 | 84.25 | **89.44** | 5.76 | 12.75 | 13.93 | 1.9 |
| | | 2 | 40 | 101.75 | 108.24 | 3.95 | 82.25 | 86.38 | 3.44 | 21.75 | 21.86 | 2.05 |
| | | 3 | 40 | 104.75 | 107.03 | 3.65 | 80.25 | 80.6 | 2.85 | 25.25 | 26.43 | 2.43 |
| | | 4 | 37 | 87.5 | 96.89 | 3.56 | 66.5 | 67.58 | 2.75 | 30 | 29.31 | 2.41 |
| | | 5 | 17 | 64.5 | 71.53 | 5.84 | 47 | 48.94 | 4.18 | 17.5 | 22.59 | 3.49 |
| | | 6 | 4 | 38 | 50.38 | 15.6 | 24.25 | 39.25 | 17.2 | 8.5 | 11.12 | 4.62 |
| | | 7 | 2 | 27.75 | 27.75 | 8.25 | 19.75 | 19.75 | 0.75 | 8 | 8 | 7.5 |
| | Unmedicated | 1 | 21 | 92.5 | 119.64 | 15.58 | 85 | 105.02 | 13.89 | 11.5 | 14.62 | 2.44 |
| | | 2 | 20 | 115.75 | 116.95 | 9.01 | 85 | 91.35 | 6.88 | 23 | 25.6 | 3.82 |
| | | 3 | 18 | 107 | 108.39 | 7.98 | 81.25 | 84.61 | 7.93 | 24.25 | 23.78 | 2 |
| | | 4 | 16 | 84.5 | 81.84 | 6.89 | 61 | 56.78 | 4.82 | 21 | 25.06 | 3.84 |
| | | 5 | 9 | 70.5 | 69.61 | 8.93 | 51 | 46.67 | 6.25 | 24 | 22.94 | 3.93 |
| | | 6 | 3 | 33 | 43.17 | 17.91 | 32 | 41 | 17.34 | 2 | 2.17 | 0.73 |
| | | 7 | 1 | 25 | 25 | | 18.5 | 18.5 | | 6.5 | 6.5 | |
| | Medicated 7d | 1 | 21 | 216.5 | **234.83/***/ ###** | 19.67 | 194.5 | **216.67/***/###** | 18.74 | 17 | 18.17 | 3.31 |
| | | 2 | 19 | 125.5 | 126.37 | 11.68 | 99.5 | 101.55 | 10.68 | 22 | 24.82 | 4.1 |
| | | 3 | 11 | 113.5 | 113 | 12.09 | 90.5 | 92.73 | 8.89 | 18.5 | 20.27 | 4.46 |
| | | 4 | 2 | 93.75 | 93.75 | 36.75 | 65.5 | 65.5 | 10 | 28.25 | 28.25 | 26.75 |
| | | 5 | 1 | 38.5 | 38.5 | | 28 | 28 | | 10.5 | 10.5 | |
| C | Controls | 1 | 28 | 96.75 | **97.11** | 7.29 | 76 | **79.16** | 5.91 | 15.25 | **17.95** | 2.11 |
| | | 2 | 28 | 98.25 | **105.45** | 5.16 | 74.25 | **82.64** | 4.18 | 20.75 | 22.8 | 2.72 |
| | | 3 | 28 | 104.5 | 104.07 | 3.51 | 74.75 | 77.96 | 3.09 | 27 | 26.11 | 1.85 |
| | | 4 | 26 | 105.5 | 102.65 | 3.07 | 71.5 | 72.75 | 2.42 | 29.75 | 29.9 | 2.3 |
| | | 5 | 13 | 85 | 85.62 | 5.26 | 69 | 65.81 | 4.66 | 16.5 | 19.81 | 3.54 |
| | Medicated 7d | 1 | 21 | 203.5 | **222.43/*** ** | 25.83 | 185.5 | **202.4/*** ** | 24.79 | 14.5 | 20.02 | 3.39 |
| | | 2 | 18 | 127 | 125.39 | 11.53 | 96.5 | 96 | 9.8 | 26.5 | 29.39 | 4.55 |
| | | 3 | 10 | 126.75 | 132.75 | 13.52 | 84.5 | 96.65 | 13.38 | 33 | 36.1 | 6.05 |
| | | 4 | 5 | 91 | 93.4 | 6.76 | 57 | 61.6 | 6.18 | 30.5 | 31.8 | 11.14 |
| | | 5 | 1 | 110 | 110 | | 44 | 44 | | 66 | 66 | |
| | Medicated 28d | 1 | 20 | 208.75 | **220.4/*** ** | 22.79 | 183.5 | **193.15/*** ** | 22.6 | 25 | **27.25/* ** | 3.62 |
| | | 2 | 18 | 126.25 | **136.14/* ** | 12.05 | 99.5 | **108.97/* ** | 10.29 | 27 | 27.17 | 4.55 |
| | | 3 | 13 | 92.5 | 91.12 | 14.03 | 60 | 65.19 | 12.07 | 17 | 25.92 | 6.21 |
| | | 4 | 6 | 102.5 | 98.08 | 16.29 | 67.5 | 70.5 | 14.19 | 20.75 | 27.58 | 12.23 |
| | | 5 | 2 | 40.75 | 40.75 | 12.75 | 28.75 | 28.75 | 2.25 | 12 | 12 | 10.5 |

| | | | | | | | | | | | | |
|---|---|---|---|---|---|---|---|---|---|---|---|---|
| | | 6 | 2 | 26.5 | 26.5 | 8.5 | 23 | 23 | 5.5 | 3.5 | 3.5 | 3 |
| REM non-suppressing medication: | | | | | | | | | | | | |
| A | Controls | 1 | 40 | 87 | 91.72 | 4.89 | 70.75 | 75.44 | 4.85 | 15.75 | 16.29 | 1.37 |
| | | 2 | 40 | 99 | **100.76** | 3.14 | 78.75 | 80.78 | 2.64 | 19.75 | 19.99 | 1.59 |
| | | 3 | 39 | 114 | 114.51 | 4.71 | 84.5 | 83.38 | 3.4 | 30.5 | 31.13 | 2.9 |
| | | 4 | 33 | 99 | 102.64 | 5.15 | 69 | 71.17 | 4.26 | 28.5 | 31.47 | 2.58 |
| | | 5 | 15 | 72.5 | 77.73 | 4.8 | 58.5 | 56.1 | 2.66 | 14 | 21.63 | 4.01 |
| | | 6 | 2 | 65.75 | 65.75 | 18.75 | 54.75 | 54.75 | 13.75 | 11 | 11 | 5 |
| | Medicated | 1 | 13 | 124.5 | 163.04 | 34.59 | 107 | 141.85 | 34.94 | 18 | 21.19 | 3.92 |
| | | 2 | 11 | 126 | **126.64/*** | 9.91 | 92 | 95.41 | 8.25 | 25 | 31.23 | 4.96 |
| | | 3 | 10 | 119 | 115.4 | 11.55 | 76.25 | 85.5 | 9.87 | 24.75 | 29.9 | 5.03 |
| | | 4 | 7 | 95 | 95.57 | 3.63 | 56.5 | 59.71 | 4 | 38 | 35.86 | 7.01 |
| | | 5 | 2 | 108.5 | 108.5 | 9.5 | 60 | 60 | 6.5 | 48.5 | 48.5 | 16 |
| | | 6 | 1 | 92 | 92 | | 68 | 68 | | 24 | 24 | |
| B | Controls | 1 | 40 | 96.25 | 103.36 | 6.65 | 84.25 | 89.44 | 5.76 | 12.75 | 13.93 | 1.9 |
| | | 2 | 40 | 101.75 | 108.24 | 3.95 | 82.25 | 86.38 | 3.44 | 21.75 | 21.86 | 2.05 |
| | | 3 | 40 | 104.75 | 107.03 | 3.65 | 80.25 | 80.6 | 2.85 | 25.25 | 26.43 | 2.43 |
| | | 4 | 37 | 87.5 | 96.89 | 3.56 | 66.5 | 67.58 | 2.75 | 30 | 29.31 | 2.41 |
| | | 5 | 17 | 64.5 | 71.53 | 5.84 | 47 | 48.94 | 4.18 | 17.5 | 22.59 | 3.49 |
| | | 6 | 4 | 38 | 50.38 | 15.6 | 24.25 | 39.25 | 17.2 | 8.5 | 11.12 | 4.62 |
| | | 7 | 2 | 27.75 | 27.75 | 8.25 | 19.75 | 19.75 | 0.75 | 8 | 8 | 7.5 |
| | Unmedicated | 1 | 17 | 104 | 126.82 | 13.1 | 88.5 | 105.59 | 11.76 | 19.5 | **21.24/*** | 2.15 |
| | | 2 | 17 | 119 | 123.35 | 9.59 | 87.5 | 95.35 | 9.13 | 29 | 28 | 3.35 |
| | | 3 | 15 | 102.5 | 101 | 4.13 | 74 | 74.37 | 4.07 | 26 | 26.63 | 2.43 |
| | | 4 | 12 | 86 | 90.46 | 8.57 | 68 | 63.62 | 3.18 | 17.75 | 26.83 | 6.47 |
| | | 5 | 5 | 88 | 80.6 | 16.85 | 54 | 46.1 | 8.19 | 39.5 | 34.5 | 9.86 |
| | | 6 | 2 | 63.75 | 63.75 | 15.25 | 56.5 | 56.5 | 12.5 | 7.25 | 7.25 | 2.75 |
| | Medicated 7d | 1 | 17 | 133 | 140.65 | 20.95 | 124 | 118 | 17.65 | 16.5 | 22.65 | 4.57 |
| | | 2 | 17 | 120.5 | 123.15 | 8.37 | 97 | 100.56 | 7.16 | 24.5 | 22.59 | 2.32 |
| | | 3 | 16 | 110 | 112.78 | 8.13 | 87.5 | 87.84 | 5.72 | 19.25 | 24.94 | 4.47 |
| | | 4 | 10 | 95.25 | 101.15 | 9.64 | 72.75 | 70.45 | 5.81 | 29 | 30.7 | 5.16 |
| | | 5 | *3* | 81.5 | 76 | 6.26 | 69 | 65.5/* | 4.8 | 10 | *10.5/*** | 1.89 |
| | | 6 | 2 | 55.75 | 55.75 | 2.25 | 40 | 40 | 8.5 | 15.75 | 15.75 | 10.75 |
| | | 7 | 1 | 48.5 | 48.5 | | 44.5 | 44.5 | | 4 | 4 | |
| C | Controls | 1 | 28 | 96.75 | **97.11** | 7.29 | 76 | 79.16 | 5.91 | 15.25 | 17.95 | 2.11 |

| | | | | | | | | | | | |
|---|---|---|---|---|---|---|---|---|---|---|---|
| | 2 | 28 | 98.25 | 105.45 | 5.16 | 74.25 | 82.64 | 4.18 | 20.75 | 22.8 | 2.72 |
| | 3 | 28 | 104.5 | 104.07 | 3.51 | 74.75 | 77.96 | 3.09 | 27 | 26.11 | 1.85 |
| | 4 | 26 | 105.5 | 102.65 | 3.07 | 71.5 | 72.75 | 2.42 | 29.75 | 29.9 | 2.3 |
| | 5 | 13 | 85 | 85.62 | 5.26 | 69 | 65.81 | 4.66 | 16.5 | 19.81 | 3.54 |
| Medicated 7d | 1 | 9 | 139 | 170.11 | 32.7 | 119.5 | 150.39 | 31.39 | 18.5 | 19.72 | 4.54 |
| | 2 | 9 | 120.5 | 123.72 | 9.85 | 98 | 90 | 7.67 | 34.5 | 33.72 | 5.21 |
| | 3 | 6 | 107.75 | 109.25 | 11.15 | 82.25 | 81.42 | 7.22 | 26.5 | 27.83 | 5.43 |
| | 4 | 4 | 77.5 | 82 | 8.88 | 62.25 | 69.5 | 9.91 | 8.5 | *12.5*/* | 4.51 |
| | 5 | 1 | 18.5 | 18.5 | | 17.5 | 17.5 | | 1 | 1 | |
| | 6 | 1 | 116.5 | 116.5 | | 52 | 52 | | 64.5 | 64.5 | |
| Medicated 28d | 1 | 9 | 152.5 | **166.89/*** | 28.44 | 105 | 134.11 | 29.51 | 21.5 | 32.78 | 8.04 |
| | 2 | 9 | 119 | 125.06 | 12.08 | 92 | 97.39 | 7.59 | 26.5 | 27.67 | 7.2 |
| | 3 | 8 | 103.75 | 107.62 | 8.42 | 74.25 | 81.44 | 7.85 | 24.25 | 26.19 | 5.08 |
| | 4 | 4 | 86.5 | 77.12 | 15.76 | 53.25 | 49.62 | 10.83 | 21.5 | 27.5 | 10.07 |
| | 5 | 2 | 81.75 | 81.75 | 18.75 | 35.75 | 35.75 | 8.75 | 46 | 46 | 10 |
| | 6 | 1 | 133 | 133 | | 62 | 62 | | 71 | 71 | |

Sleep cycle durations median, mean and standard error (SE) split by non-REM and REM parts defined according to Feinberg & Floyd (1979). Note that the number of available sleep cycles (N) drop sharply after 3rd or 4th cycle. Medicated patients in all datasets have extended first and second sleep cycles. This effect seemed to be driven by extension of non-REM periods of the sleep cycle. REM sleep was extended after prolonged sleep cycles, suggesting higher REM pressure, in the long-term medication groups (i.e. Dataset A Medication, Dataset C Medicated 28d). Effects seemed to be driven by patients that received REM suppressing medication as above effects were lessened under medication not known to suppress REM. Different symbols are used for indicating statistical comparisons (two-tailed paired and unpaired two sample *t*-tests with assumptions for unequal variance) within the datasets that are significant (highlighted in bold, red higher, blue lower): differences to Controls use asterisks (*, $p<.05$; **, $p<.01$; ***, $p<.001$), within patients for their follow-ups (Dataset B Unmedicated vs. Medicated 7d; Dataset C Medicated 7d vs. Medicated 28d) use hashes (###, $p<.001$). Note that some statistics were run on very low <10 sleep cycles in one of the groups and were thus not interpreted.

**Table S5**

Sleep parameters of all three datasets combined (mean ± SE).

| | Combined datasets | |
|---|---|---|
| | Controls | Medicated |
| | Sleep architecture | |
| N1[%] | **10.8 ± 0.523** | **15.1 ± 0.699*****|
| N2[%] | 46.6 ± 0.79 | 49 ± 1.04 |
| SWS[%] | **17 ± 0.76** | **14.2 ± 0.891**** |
| Non-REM[%] | 63.6 ± 0.7 | 63.3 ± 1 |
| REM [%] | **17.9 ± 0.507** | **13.1 ± 0.737**** |
| WASO [%] | 7.45 ± 0.619 | 7.99 ± 0.57 |
| TST [min] | 466 ± 2.04 | 465 ± 1.76 |
| Sleep onset [min] | 21 ± 1.26 | 24.4 ± 1.39 |
| SWS onset [min] | **19.7 ± 1.36** | **27.6 ± 2.42**** |
| REM onset [min] | **81 ± 3.27** | **179 ± 9.9***** |
| | Sleep spindles | |
| Density [/epoch] | 2.25 ± 0.039 | 2.16 ± 0.046 |
| Count | 1292 ± 28.4 | 1223 ± 36.9 |
| Amplitude [µV] | 28.7 ± 0.871 | 28.3 ± 0.799 |
| Frequency [Hz] | 13.2 ± 0.05 | 13.1 ± 0.06 |
| Duration [s] | 0.78 ± 0.005 | 0.783 ± 0.005 |
| | Slow waves | |
| Density [/epoch] | 1.47 ± 0.035 | 1.46 ± 0.039 |
| Count | 845 ± 24.5 | 830 ± 29.8 |
| Amplitude [µV] | **158 ± 3.18** | **141 ± 3.37***** |
| Frequency [Hz] | **0.811 ± 0.01** | **0.789 ± 0.01**** |
| Duration [s] | **1.24 ± 0.008** | **1.27 ± 0.009**** |
| | SW-spindles | |
| Count | 96.1 ± 4.72 | 86.1 ± 4.7 |
| Mean delay [s] | 0.551 ± 0.006 | 0.567 ± 0.006 |
| Delay dispersion [sd] | **0.221 ± 0.004** | **0.244 ± 0.004***** |

Controls (n = 108), Medicated patients (n = 108). Differences between controls and medicated patients in use the following: **, *p*<.01; ***, *p*<.001.

**Table S6**

Behavioral results of finger tapping test of Dataset A and Dataset B combined

| | Controls (comb) | Medicated (comb) |
|---|---|---|
| Baseline | **6.95 ± 0.395 (1 rm)** | **5.66 ± 0.488 *** |
| Training | 1.51 ± 0.158 (2 rm) | 1.27 ± 0.126 (1 m) |
| Consolidation | **0.097 ± 0.023** | **-0.07 ± 0.036 $^{***}$(2 rm)** |

Results are reported after removal of outliers 3sd from the overall mean Differences between controls and medicated patients in use the following: *, p <0.05; ***, p<0.001. 1/2 rm = one/two outlier(s) more than 3sd from the mean removed

**Table S7**

Medication of patients

| Participant | Dataset | Medication type |
|---|---|---|
| 1 | Dataset A | Sertraline |
| 2 | Dataset A | Mirtazapine |
| 3 | Dataset A | Venlafaxine, Quetiapine |
| 4 | Dataset A | Amitriptyline, Amlodipine |
| 5 | Dataset A | *Medication type unknown* |
| 6 | Dataset A | Mirtazapine, Trimipramine, Sertraline |
| 7 | Dataset A | Mirtazapine, Venlafaxine, Lorazepam |
| 8 | Dataset A | Venlafaxine, Olanzapine, Ramipril, Propranolol |
| 9 | Dataset A | *Medication type unknown* |
| 10 | Dataset A | *Medication type unknown* |
| 11 | Dataset A | Duloxetine, Amitriptyline, Escitalopram |
| 12 | Dataset A | Escitalopram, Trimipramine |
| 13 | Dataset A | Venlafaxine |
| 14 | Dataset A | Quetiapine, Amitrityline, Lorazepam |
| 15 | Dataset A | Mirtazapine |
| 16 | Dataset A | Duloxetine, Trimipramine, Lithium, Lorazepam, L-Thyroxin |
| 17 | Dataset A | Citalopram, Amitriptyline, Zopiclon, Bisohexal, Atorvastatin, L-Thyroxin |
| 18 | Dataset A | Mirtazapine |
| 19 | Dataset A | Sertraline, Zopiclon, Bisohexal, Atorvastatin, L-Thyroxin |
| 20 | Dataset A | *Medication type unknown* |
| 21 | Dataset A | Quetiapine, Escitalopram, Venlafaxine, Lamotrigine, Olanzapin |
| 22 | Dataset A | Lithium, Venlafaxine, Escitalopram, Pregabalin |
| 23 | Dataset A | Escitalopram, Lorazepam, Pantoprazol, L-Thyroxin |
| 24 | Dataset A | *Medication type unknown* |
| 25 | Dataset A | Trimipramine, Sulpiride, Lithium |
| 26 | Dataset A | Quetiapine, Escitalopram, L-Thyroxin |
| 27 | Dataset A | Trimipramine, Escitalopram, Lamotrigin |

| | | |
|---|---|---|
| 28 | Dataset A | Venlafaxine, Escitalopram, Mirtazapine, Thyronajod |
| 29 | Dataset A | Citalopram, Trimipramine, Lamotrigin |
| 30 | Dataset A | Quetiapine, Mirtazapine, Venlafaxine, Metroprolol, Enahexal |
| 31 | Dataset A | *Medication type unknown* |
| 32 | Dataset A | Doxepine |
| 33 | Dataset A | Duloxetine, Pregabaline, Enahexal, Metohexal, Pantoprazol |
| 34 | Dataset A | Mirtazapine, Trimipramine, Sertraline |
| 35 | Dataset A | Clomipramine |
| 36 | Dataset A | Mirtazapine |
| 37 | Dataset A | *Medication type unknown* |
| 38 | Dataset A | Mirtazapine, Duloxetine |
| 39 | Dataset A | Venlafaxine, Lorazepam, Zopiclon |
| 40 | Dataset A | Trimipramine, Sertraline |
| 1 | Dataset B | Citalopram |
| 2 | Dataset B | Duloxetine |
| 3 | Dataset B | Venlafaxine |
| 4 | Dataset B | Citalopram |
| 5 | Dataset B | Venlafaxine |
| 6 | Dataset B | Venlafaxine |
| 7 | Dataset B | Trimipramine |
| 8 | Dataset B | Citalopram |
| 9 | Dataset B | Bupropion |
| 10 | Dataset B | Citalopram |
| 11 | Dataset B | Citalopram |
| 12 | Dataset B | Citalopram |
| 13 | Dataset B | Escitalopram |
| 14 | Dataset B | Escitalopram |
| 15 | Dataset B | Amitriptyline |
| 16 | Dataset B | Paroxetine |
| 17 | Dataset B | Trimipramine |
| 18 | Dataset B | Duloxetine |
| 19 | Dataset B | Amitriptylineoxide |
| 20 | Dataset B | Venlafaxine |

| | | |
|---|---|---|
| 21 | Dataset B | Escitalopram |
| 22 | Dataset B | Trimipramine |
| 23 | Dataset B | Mirtazapine |
| 24 | Dataset B | Escitalopram |
| 25 | Dataset B | Escitalopram |
| 26 | Dataset B | Mirtazapine |
| 27 | Dataset B | Trimipramine |
| 28 | Dataset B | Mirtazapine |
| 29 | Dataset B | Trimipramine |
| 30 | Dataset B | Trimipramine |
| 31 | Dataset B | Bupropion |
| 32 | Dataset B | Trimipramine |
| 33 | Dataset B | Sertraline |
| 34 | Dataset B | Bupropion |
| 35 | Dataset B | Bupropion |
| 36 | Dataset B | Mirtazapine |
| 37 | Dataset B | Bupropion |
| 38 | Dataset B | Bupropion |
| 39 | Dataset B | Bupropion |
| 40 | Dataset B | Mirtazapine |
| 1 | Dataset C | Mirtazapine |
| 2 | Dataset C | Venlafaxine |
| 3 | Dataset C | Reboxetine |
| 4 | Dataset C | Venlafaxine |
| 5 | Dataset C | Venlafaxine |
| 6 | Dataset C | Venlafaxine |
| 7 | Dataset C | Cymbalta, Elontril |
| 8 | Dataset C | Duloxetine, Trimipramine |
| 9 | Dataset C | Equilibrine, Stangyl, Tavor |
| 10 | Dataset C | Sertraline, Lamotrigine, Olanzapine |
| 11 | Dataset C | Doxepine |
| 12 | Dataset C | Elontril |
| 13 | Dataset C | Trimipramine |

| | | |
|---|---|---|
| 14 | Dataset C | Venlafaxine |
| 15 | Dataset C | Venlafaxine |
| 16 | Dataset C | Venlafaxine, Trimipramine, Quetiapine |
| 17 | Dataset C | Citalopram |
| 18 | Dataset C | Amitriptyline, Amitriptylinoxid |
| 19 | Dataset C | Venlafaxine |
| 20 | Dataset C | Equilibrine, Movicol |
| 21 | Dataset C | Stangyl |
| 22 | Dataset C | Mirtazapine |
| 23 | Dataset C | Paroxetine, Anafranil, Seroquel, Musaril, Piroxicam, ASS, Allopurinol, Sortis |
| 24 | Dataset C | Elontril, Trevilor, Lithium |
| 25 | Dataset C | Trimipramine, Saroten |
| 26 | Dataset C | Duloxetine, Lorazepam, Lthyron, Trazodon |
| 27 | Dataset C | Venlafaxine, Mirtazapine, LTG, Lorazepam, Thyronajod |
| 28 | Dataset C | Venlafaxine, Mirtazapine, Lorazepam, Gabapentin |
| 29 | Dataset C | Elontril, Zyprexa |
| 30 | Dataset C | Venlafaxine |
